# Discovering plasticity rules that organize and maintain neural circuits

**DOI:** 10.1101/2024.11.18.623688

**Authors:** David Bell, Alison Duffy, Adrienne Fairhall

## Abstract

Intrinsic dynamics within the brain can accelerate learning by providing a prior scaffolding for dynamics aligned with task objectives. Such intrinsic dynamics should self-organize and self-sustain in the face of fluctuating inputs and biological noise, including synaptic turnover and cell death. An example of such dynamics is the formation of sequences, a ubiquitous motif in neural activity. The sequence-generating circuit in zebra finch HVC provides a reliable timing scaffold for motor output in song and demonstrates a remarkable capacity for unsupervised recovery following perturbation. Inspired by HVC, we seek a local plasticity rule capable of organizing and maintaining sequence-generating dynamics despite continual network perturbations. We adopt a meta-learning approach introduced by Confavreux et al, which parameterizes a learning rule using basis functions constructed from pre- and postsynaptic activity and synapse size, with tunable time constants. Candidate rules are simulated within initially random networks, and their fitness is evaluated according to a loss function that measures the fidelity with which the resulting dynamics encode time. We use this approach to introduce biological noise, forcing meta-learning to find robust solutions. We first show that, in the absence of perturbation, meta-learning identifies a temporally asymmetric generalization of Oja’s rule that reliably organizes sparse sequential activity. When synaptic turnover is introduced, the learned rule incorporates an additional form of homeostasis, better maintaining sequential dynamics relative to other previously proposed rules. Additionally, inspired by recent findings demonstrating plasticity in synapses from inhibitory interneurons in HVC, we explore the role of inhibitory plasticity in sequence-generating circuits. We find that learned plasticity adjusts both excitation and inhibition in response to manipulations, outperforming rules applied only to excitatory connections. We demonstrate how plasticity acting on both excitatory and inhibitory synapses can better shape excitatory cell dynamics to scaffold timing representations.

## 1 Introduction and related work

How computational structures are organized and maintained within the brain is a central question within neuroscience. While feedback is clearly essential for learning, self-organization of neural circuits can unfold without feedback, e.g. during development. Brains have evolved specific cell types with nonrandom spatial organization, plasticity rules, and connectivity that likely introduce a strong set of inductive biases on the information processing they perform. How might organization of useful circuit dynamics be established and maintained throughout life without the need for feedback? Recent work suggests self-organized computations, once established, can accelerate learning and improve performance when experience is limited: Nicola and Clopath [1] demonstrated that a stable high dimensional time signal could improve a network’s performance on sequential motor tasks (Fig. 1a). Najarro and Risi [2] learned Hebbian plasticity that orchestrated spontaneous walking behavior in quadruped agents; similar work has shown architectural priors increase the sample efficiency and generalization of RL approaches to locomotion [3, 4]. Additionally, in RL settings, supplying agents with a time input permits them to adopt time-dependent policies [5]. The ability of computational primitives, such as timing representations, to self-organize is challenged by the shifting structure of neural circuits. Synaptic loss, synaptogenesis, cell death, and neurogenesis pose challenges for all learning algorithms, but particularly for self-organization which must be based solely on local information rather than global task performance.

**Figure 1:**
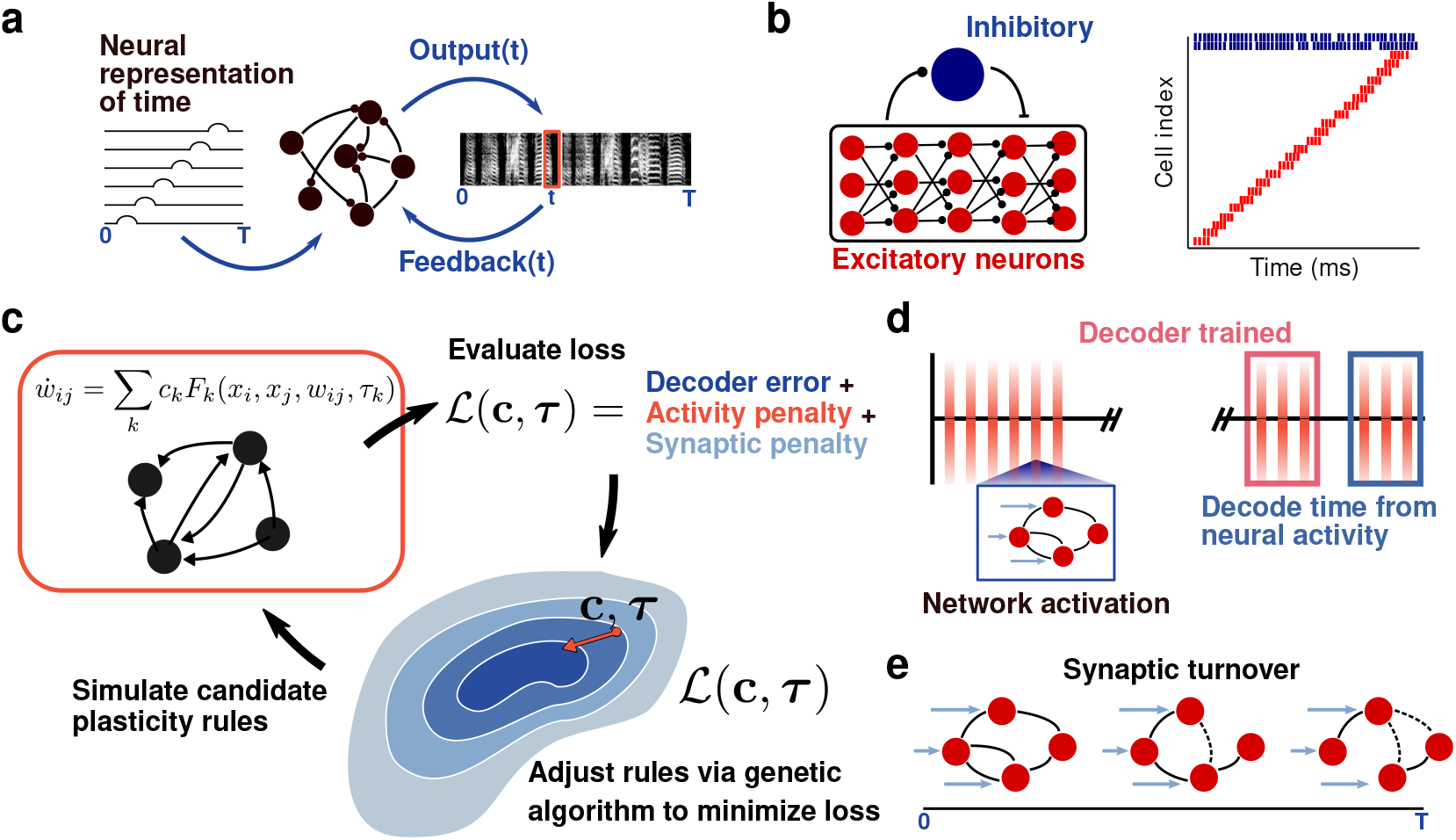
Meta-learning approach to discovering plasticity rules that organize sequences. **(a)** In zebra finch song learning, a neural representation of time (left) in HVC simplifies the sequential motor learning task of producing the correct spectral output. **(b)** Putative network structure of zebra finch HVC: a feed-forward, excitatory network with recurrent inhibition (left). HVC excitatory neurons (red) fire sparsely in time while interneurons (blue) fire tonically (right). **(c)** Strategy for learning plasticity underlying sequence organization: candidate plasticity rules, parameterized by a set of coefficients and time constants, are simulated. A loss function is evaluated on the resulting dynamics, and new candidate rules are generated. **(d)** Test procedure for representation of time. Networks are activated 400 times (blue bars). From the final 50 activations, 6 are chosen to train a decoder and 6 to test the representation by decoding time from neural activity. **(e)** The robustness of discovered plasticity rules can be tested by introducing synaptic turnover, in which synapses are stochastically removed and added, into simulations.

Here, we aim to find plasticity rules that self-organize and maintain one useful computational primitive: sparse, sequential activity. Such activity is widely seen in many areas of the brain including hippocampus [6], cortex [7], and basal ganglia [8]. In the songbird zebra finch, area HVC (used as a proper noun), a cortical-like region, displays sequential activity representing time [9], reducing the problem of motor learning to driving the correct motor neuron at the correct moment [10]. Extensive literature has explored how such sequence-generating circuits could emerge in the absence of feedback [11–17], but has largely focused on either how these structures organize or how they self-maintain, using guessed plasticity rules, and neglecting the effects of ongoing synaptic noise. Further, previous work on sequence organization within HVC has focused on plasticity between excitatory (E) neurons. Recent experimental findings show unsupervised recovery of HVC dynamics is accompanied by changes in both E→E and also inhibitory-to-excitatory (I→E) synaptic strength [18].

Here, instead of imposing a guessed rule, we ask which self-supervised plasticity rules can organize and maintain sequential dynamics within a network. We employ meta-learning, a supervised method to learn learning rules [19–23], in order to discover rules that self-organize a sequence. Our approach stems from a rich history of learning local plasticity rules, including rules that extract representations from data [24], enhance artificial agent performance on familiarity and navigation tasks [2, 25], and explore biologically-plausible replacements or complements to backpropagation [19, 20, 26]. In this study, we parameterize the space of plasticity rules with a basis of activity- and synapse size-dependent terms. The set of coefficients weighting these terms and associated time constants are adjusted to minimize a loss function. Inspired by the HVC context, we pose the loss in terms of the accuracy with which the time since an initial network input can be decoded from the circuit dynamics. We first consider E→E rules alone, and then add I→E and E→I plasticity. We then introduce perturbations to the circuit and investigate which learned plasticity rules promote circuit stability. We find that meta-learned rules for self-organized sequence generation and maintenance contain distinct forms of spike-timing dependent plasticity, homeostasis, and network bounds, which outperform previously proposed sequence-organizing rules in the presence of noise and that plasticity on reciprocal connectivity to inhibitory neurons confers additional stability on the network dynamics. Our main contribution is the exploration of unsupervised and unrewarded plasticity via meta-learning that organizes and maintains a specific and biologically relevant computational motif, a sequence.

### 1.1 Background on zebra finch physiology

In the zebra finch, nucleus HVC contains excitatory neurons that fire sparsely (typically in one burst of spikes) during song and are purportedly arranged in a feed-forward structure (Fig. 1b) [27]. A subset of these cells, known as HVC_(RA)_ neurons, project to downstream nucleus RA (robust nucleus of the archistriatum), which in turn projects to vocal neurons of the syrinx and to the brainstem, which regulates respiration [9]. HVC receives excitatory projections from nucleus Uva, which controls the onset of song syllables [28] and provides input for the duration of song [29]. HVC_(RA)_ neurons inhibit each other disynaptically via a population of inhibitory interneurons [30]. Remarkably, singing behavior can persist when the nucleus is transected [31], demonstrating its resilience.

## 2 Results

### 2.1 Learning biologically plausible plasticity on E→E synapses that organizes a sequence-generating circuit

We first learned a plasticity rule on E→E synapses that organizes a randomly connected network into a sequence-generating circuit in the absence of any perturbation. We initially constrained plasticity to E→E synapses to compare with previously proposed rules, which have largely only considered E→E plasticity. We did not include rules that imposed hard bounds on the size of individual synapses or on the collective strength of all synapses onto a neuron, as we aimed to learn plasticity rules that could permit flexible rescaling of connections in response to perturbations, as has been observed experimentally [18].

We use a network of 25 E and 8 I threshold-linear neurons. Each neuron fired according to *x*_*j*_(*t*) = [*V*_*j*_(*t*) − *b*]^+^, where *V*_*j*_(*t*) evolves via 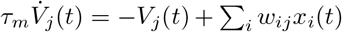.Here, *w*_*ij*_ is the weight of the synapse *i* → *j, τ*_*m*_ is the membrane time constant, and *b* is the bias. Initial connectivity (Fig. 2d) was random, but contained no I→I connectivity as is the case in HVC [30, 32] (see Supp. Sec. 1 for all network model details).

**Figure 2:**
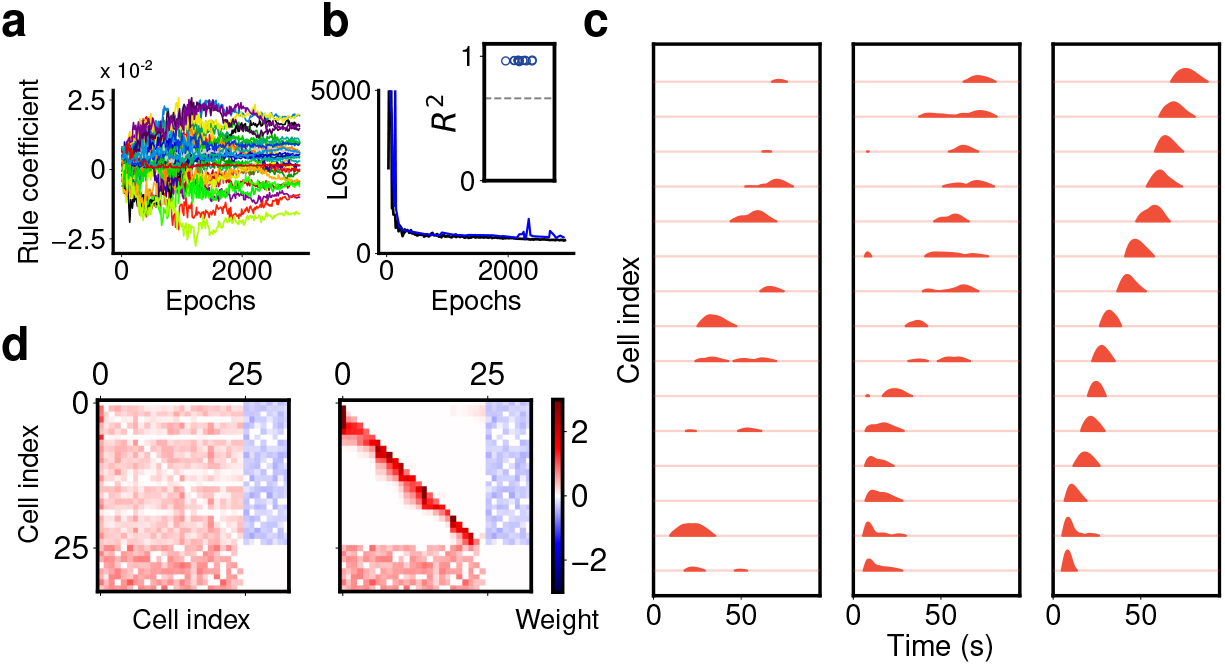
Meta-learning discovers unsupervised local plasticity rules that organize sequential activity. **(a)** Evolution of coefficients and time constants during meta-learning. **(b)** Training loss (black) and test loss (blue) during meta-learning. Inset: median decoding accuracy for 14 learned rules across 100 networks (blue points) compared to no plasticity (dashed gray line). **(c)** Meta-learned plasticity rules generate sequence dynamics. Network activity during the first activation (left), 200^*th*^ (middle), and 400^*th*^ (right), sorted by ordering of mean firing time of the final activation (425^*th*^). Note: plasticity rule is held fixed during simulation. **(d)** Weight matrices at activation 1 and 400, sorted based on mean firing time of the final activation. Connectivity between E cells organizes into a feedforward structure.

#### 2.1.1 Meta-learning procedure

We adopt a meta-learning approach pioneered by Bengio et al. [19] and extended by Confavreux et al [33]. We parameterize a set of plasticity rules with coefficients *c*_*k*_ and time constants *τ*_*k*_, such that individual synapses, *w*_*ij*_, evolve according to

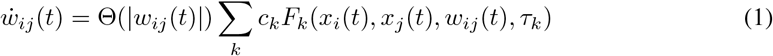

where *x*_*i*_ and *x*_*j*_ are pre- and postsynaptic activities, respectively, Θ is the Heaviside function, and *F*_*k*_ is the *k*^*th*^ term in the plasticity rule (see Fig. 3a or Supp. Sec. 2 for all terms). The basis includes terms that filter pre- and postsynaptic activities with decaying exponentials (denoted e.g. 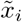) as these terms convey information about the durations of activations and relative ordering, not just their instantaneous rates. Synapses evolving under Eq. 1 were bounded so that they obeyed Dale’s law.

**Figure 3:**
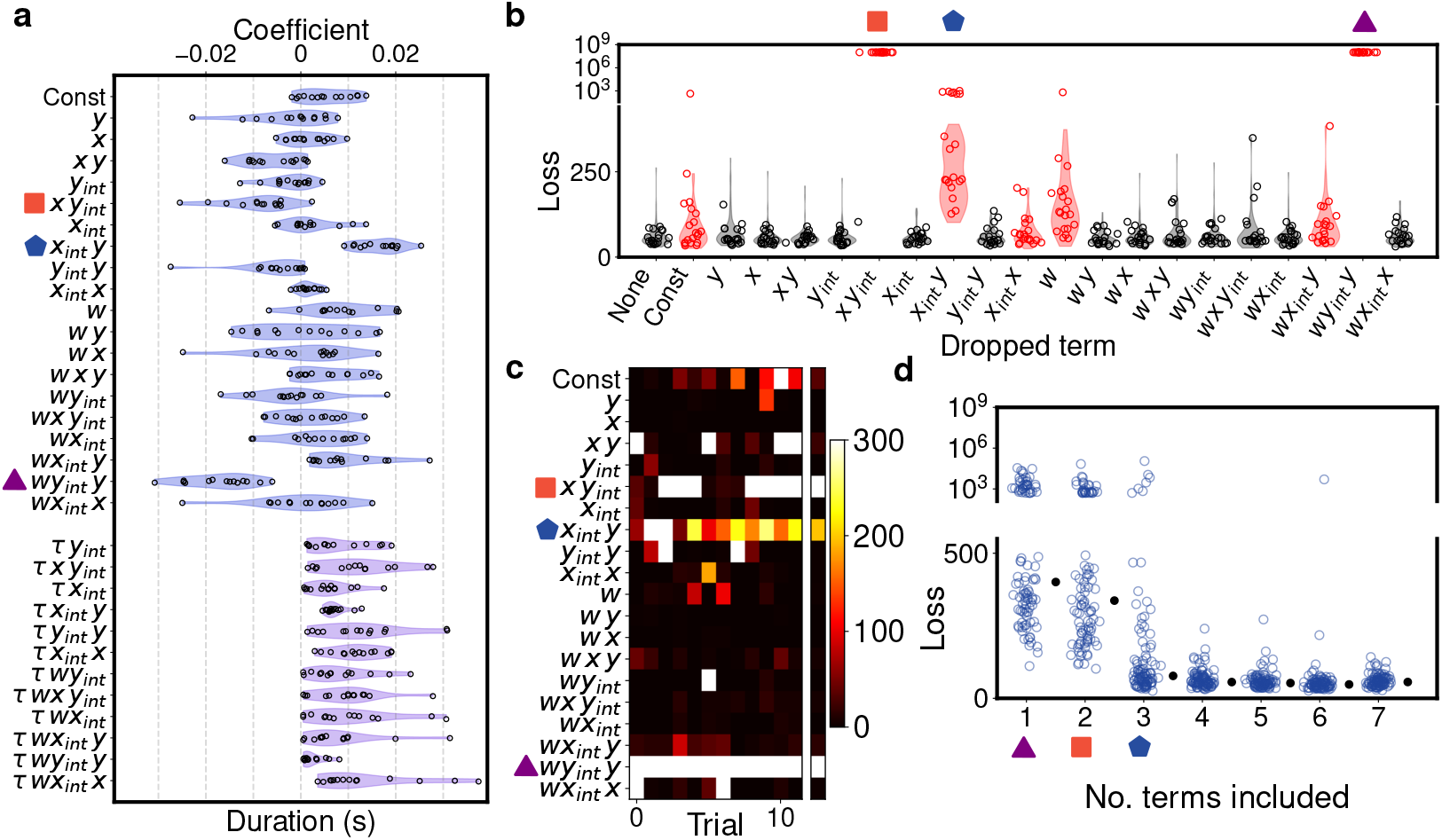
Perturbing learned plasticity rules reveals dependence on temporally asymmetric Hebbian learning and a postsynaptic activity bound. **(a)** Distributions of coefficients (blue) and time constants (purple) of basis terms across N=14 training instances (individual instances in black). The basis set consists of functions of pre and post-synaptic neural activity and synaptic weights. *x* = activity of pre-synaptic neuron; *y* = activity of post-synaptic neuron; *w* = weight of synapse; 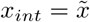(*x* filtered with *τ*). **(b)** Performance loss when individual coefficients are set to zero. **(c)** Difference in median loss between full learned solutions run on 100 test networks and solutions with one term dropped. Separate column denotes median loss for terms across all trials. **(d)** Progressive refitting of model in order of impact on median loss with and without term. Symbols indicate the term added at each refitting. Losses for 100 test networks shown (blue); medians shown in black.

We next define a loss function that evaluates the quality of the sequential dynamics organized under a chosen {*c*_*k*_, *τ*_*k*_} and attempt to minimize it using an evolutionary strategy, Covariance Matrix Adaptation (CMA-ES) [34]. We use CMA-ES to sample from the space of possible {*c*_*k*_, *τ*_*k*_} and evaluate the loss at each point by simulating 10 randomly initialized networks under the given rule and evaluating the resultant dynamics at the end of the simulation. Each simulation is divided into 400 activations of 110 ms. At *t* = 10 ms of each activation, a single fixed neuron is driven by a strong kick of excitation [17]. Following this, all other neurons in the network receive Poisson distributed input for a period of 65 ms. A fraction of this Poisson input is held fixed from trial to trial, mirroring input to HVC from the nucleus Uva, which likely does not provide a fully stochastic signal to the downstream area, [28]. The total loss for a given rule is simply the sum of losses across each of the 10 networks.

#### 2.1.2 Loss function

To learn sequences, we define a loss function based on three principles: (1) elapsed time since initial network activation should be readily decoded from network activity, (2) total network activity should be sparse, and (3) total synaptic change in the network should be minimized. The latter two principles impose the assumption that effective plasticity does not require excess network activity or synaptic change to stabilize network function. Our loss is

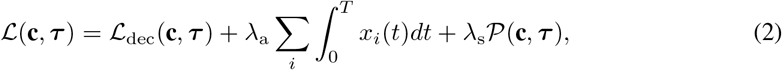

where

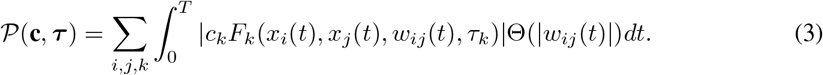

𝒫 (**c, *τ***) penalizes the all synaptic changes due to each component of **F**. This *L*_1_-like penalty on **F** penalizes each term not by the size of the term’s coefficient, *c*_*k*_, but by the quantity of synaptic change it evokes. While penalizing the magnitude of *c*_*k*_ is standard [20, 33], we take this approach to compare different terms on a common scale, as each component of **F** has differing dependence on *x*_*i*_, *x*_*j*_, and *w*_*ij*_. *λ*_a_ and *λ*_s_ are positive constants weighting the activity and synaptic change penalties.

To determine the decoder loss ℒ_dec_(**c, *τ***), the activity of networks was sampled at 500 time points of six activations of the network. From these, a linear decoder was constructed and used to decode the activity at 200 time points of six subsequent activations. Note that since the initial kick of excitation was time-locked to beginning of each activation, constructing a decoder to read out time elapsed since the beginning of the activation was equivalent to decoding time elapsed since the initial excitatory kick was presented.

#### 2.1.3 Learned E→E plasticity induces sequences using temporally asymmetric Hebbian learning and a peak postsynaptic activity bound

Meta-learning reliably found plasticity rules that organized random E synaptic connectivity into feed-forward structures that generated sequences when activated (Fig. 2d-e). The organized structures were not grouped into links, as in a synfire chain, but were better described by a kernel in which the strength of a synapse between two cells depended on the lag between their mean firing times (Fig. 2d, right). The set of rules discovered by meta-learning are not sparse in the space of {*c*_*k*_, *τ*_*k*_} (Fig. 3a). To test whether learned plasticity rules truly required all terms with nonzero coefficients, we compared ‘dropout’ variants of the discovered rule, in which one coefficient within {*c*_*k*_} was set to zero, to its unaltered form. We computed the loss of these variants on test networks to determine if a term’s absence impacted the loss (Fig. 3b). Computing the change in the median loss of networks organized by the learned solution and a dropout variant across 14 discovered rules revealed reliable trends in the importance of various terms (Fig. 3c). We refit these terms progressively in order of the impact on loss across all training runs and found a sharp elbow at 3 terms: a temporally asymmetric Hebbian learning term, 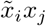 (blue pentagon in Fig. 3), its complement 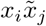 (orange square), and a term second order in the postsynaptic activity multiplied by the synapse size, 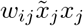 (purple triangle). The first term was consistently learned with a positive coefficient while the latter two were always almost negative (Fig. 3a), rendering the effective learning rule

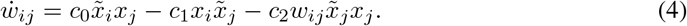

where all {*c*_*i*_} are non-negative and time constants are different for each term. The 3 most important terms can be interpreted as a temporally asymmetric generalization of Oja’s rule in that the Hebbian learning term, *x*_*i*_*x*_*j*_, is replaced by the first two terms in Eq. 4, which depend on the relative timing of *x*_*i*_ and *x*_*j*_. The time constant of the third term was on average short (∼1 ms), making this term akin to 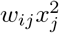, the normalizing term of Oja’s rule.

### 2.2 Biological noise alters the learned plasticity rules

We next asked whether ongoing disruptions to network structure alter which plasticity rules are meta-learned. To explore this, we introduced synaptic turnover to the simulation phase of the meta-learning loop. Synaptic turnover is a stochastic process by which existing synapses disappear and new, small synapses emerge (Fig. 1e). Prior to each network activation, all connections were updated according to

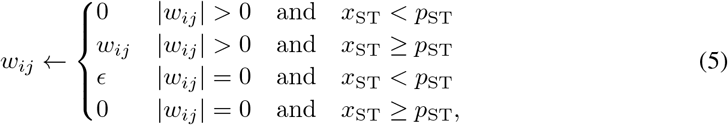

where *x*_*ST*_ ∼ *U* [0, 1], *p*_*ST*_ is the probability of single synapse turnover per activation, and *ϵ* is a small positive (negative) constant if the presynaptic cell is excitatory (inhibitory). Since plasticity is unable to act on connections of size 0 (see Eq. 1), synaptic turnover determines the set of synapses available to the plasticity rule. To ensure learned rules were robust to a spectrum of rates of synaptic turnover, only half the networks used to evaluate the batch loss underwent this process.

We found that meta-learned rules were able to organize persistent representations of time despite synaptic turnover, with performance near that of rules learned on unperturbed networks. Our term-sensitivity analysis (Fig 4a) showed that solutions again heavily depended on temporally asymmetric Hebbian learning, i.e. 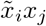 (light blue hexagon), and the bound on postsynaptic activity, 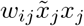 (purple triangle); however, we frequently found dependence on two additional terms: one that constantly strengthened all synapses (dark blue pentagon) and an activity bound independent of synapse size (orange square). Refitting the plasticity rule in order of impact on loss demonstrated that these 4 terms recapitulated most of the success of the learned solutions. The effective rule may be written as

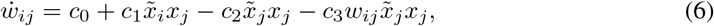

where again all {*c*_*i*_} are non-negative. Synapses for which postsynaptic activity, *x*_*j*_, remains chronically small will be potentiated by *c*_0_; however, adding this potentiating term also necessitates an activity bound that does not scale with synapse size, such as the term with prefactor *c*_2_ in Eq. 6. To understand this impact of this term, consider a synapse between two neurons whose typical firing times are far apart, i.e. 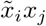 is nearly zero, the fixed point under this rule is

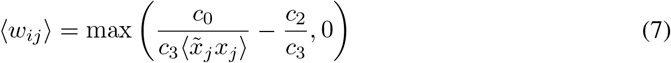

if 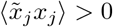,where ⟨·⟩ denotes the time average. Thus, a large enough choice of *c*_2_ prevents every synapse in the network from growing, enforcing sparsity and decreasing the risk of a neuron changing its firing time upon loss of its original inputs. We additionally note that Eq. 6 does not contain 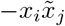,which appeared in the reduced rule learned in the absence of synaptic turnover (Eq. 4). This may be because the roles of 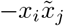 and 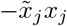 are partially redundant: both terms can suppress synapses that run counter to the sequential dynamics in the network. When constant potentiation of all synapses occurs, the term in 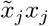 is preferable as it offsets constant potentiation of all synapses. In the unperturbed context, where constant synaptic growth is unnecessary, the term in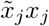 is problematic in that it can set all afferent synapses to a driven neuron to zero (whereas 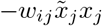 cannot). Thus, the term in 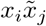 becomes preferable in the unperturbed context.

**Figure 4:**
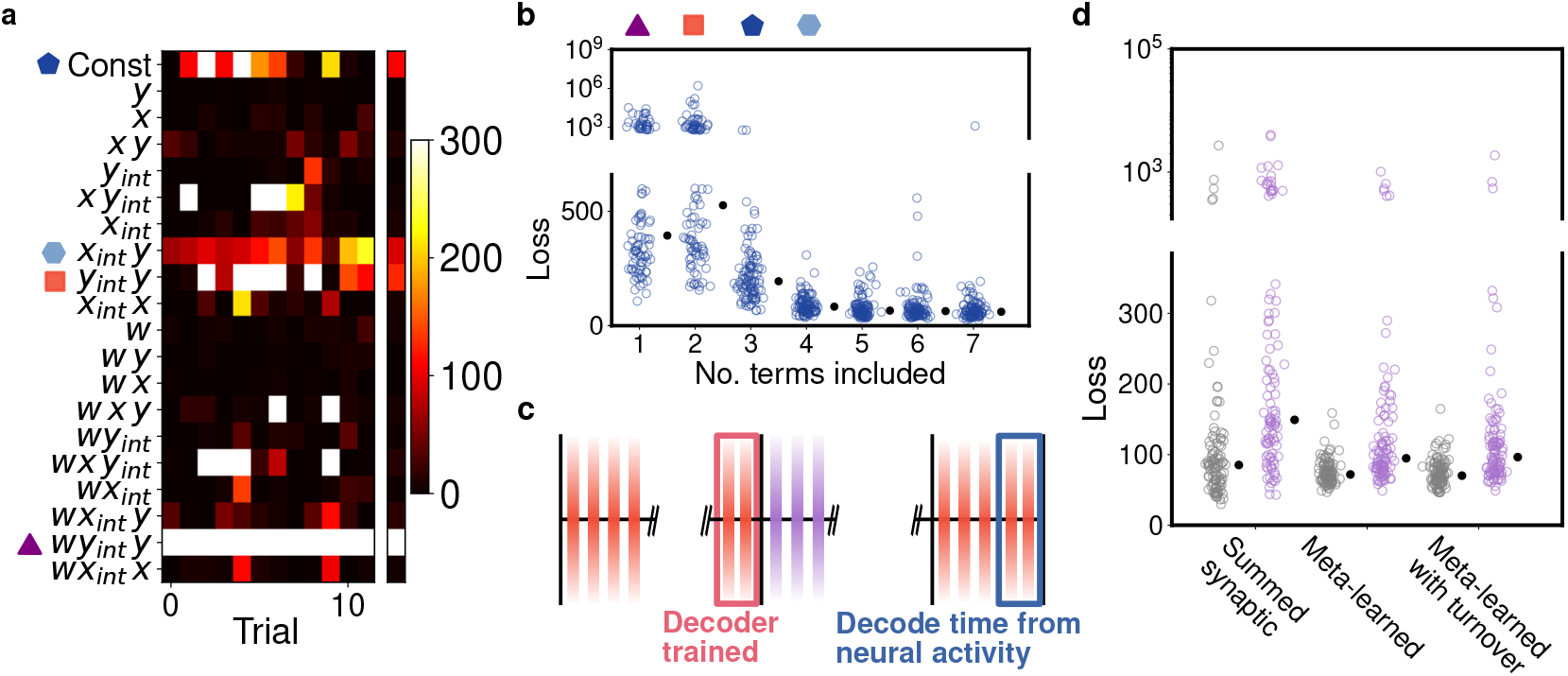
Discovered rules organized dense feedforward structures. **(a)** Median impact of loss when coefficient of given plasticity is set to zero for a given learned solution. **(b)** Refitting terms in order of median impact on loss shows sequence generation in synaptic turnover context is well captured by 4 terms. **(c)** Task structure for comparison between different rules. Rules are given 400 activations to organize dynamics. At the end, the decoder is constructed. Networks then undergo synaptic perturbation for 150 activations. Finally, the decoder attempts to decode time from the resulting dynamics to determine the loss. **(d)** Comparison between multiplicative Hebbian learning and summed synaptic bound, meta-learned without synaptic turnover, and meta-learned with synaptic turnover on time encoding task illustrated in (c). Trials shown in grey do not include synaptic turnover; light purple include synaptic turnover. Black dots indicate median values for each condition.

#### 2.2.1 Comparison to existing models of sequence formation

Do these discovered learning rules more robustly encode timing than previously proposed rules when the circuit is disrupted with biologically relevant noise? We hypothesized this would be true given that discovered rules do not impose hard bounds on the size of single synapses, total synaptic strength onto a neuron, nor number of synapses, as other models of sequence formation have [11–13, 17]. The absence of these constraints permits compensatory rescaling of synapses in response to disruptions. We compared meta-learned rules trained with and without synaptic turnover to a previously proposed sequence learning rule that used multiplicative asymmetric Hebbian learning, a single synapse bound, and bounds on the total strength of synapses onto and out of individual neurons [11]. Each rule was applied to 100 test networks for 400 activations without disruption. Following this, a decoder was constructed to read out time from the neural activity, and then synapses in the networks were turned over during an additional 150 activations, after which the loss was evaluated using the constructed decoder (Fig. 4c). We compared the best versions of each rule when synapses were turned over (with probability *p*_*ST*_ = 0.00072) at each activation, equivalent to a 90% probability of individual synapse survival during the disruption period (Fig. 4d, light purple points), and when they were not turned over (Fig. 4d, grey points; see Supp. Sec. 4). We used the Kruskal-Wallis *H* test to test for equality of medians. We found that meta-learned rules trained with and without synaptic turnover outperformed the rule based based on rigid synapse constraints (*p* = 2.5 × 10^−8^, Cohen’s *d* = −0.45 and *p* = 7.2 × 10^−7^, Cohen’s *d* = −0.44, respectively), while in the absence of perturbation, medians were not distinct after 4-fold Bonferroni correction (*p* = 0.016, Cohen’s *d* = −0.29 and *p* = 0.029, Cohen’s *d* = −0.30, respectively). When we studied the connectivity structure of networks organized by meta-learned rules, we found that the discovered rule generated denser feedforward connectivity in comparison to other plasticity rules with alternative forms of Hebbian learning and heterosynaptic competition (see Supp. Sec. 5).

### 2.3 Including inhibitory plasticity

Meta-learning allows the exploration of multiple plasticity rules operating on distinct sets of synapses within the same circuit, as might arise if there are multiple cell types [35, 36]. In particular, I→E plasticity has been the focus of much recent work [37–42]. The interaction of many plasticity rules is challenging to analyze theoretically, but meta-learning allows exploration of these interactions [43, 44]. Recent evidence suggests there may be multiple forms of plasticity within sequence-generating circuits: Wang et al. [18] increased the intrinsic excitability of *in vivo* HVC_(RA)_ neurons and found that the strength of both E→I and I→E connections could dynamically shift in response. Targeted cells received increased total inhibitory synaptic strength and decreased excitatory strength.

To understand the role of this additional plasticity, we meta-learned plasticity on all sets of synapses within the circuit (Fig. 5a) both with and without synaptic turnover. Specifically, we attempted to learn three independent plasticity rules that operated on three distinct groups of synapses (E→E, E→I, and I→E; see Supplementary Section 6).

**Figure 5:**
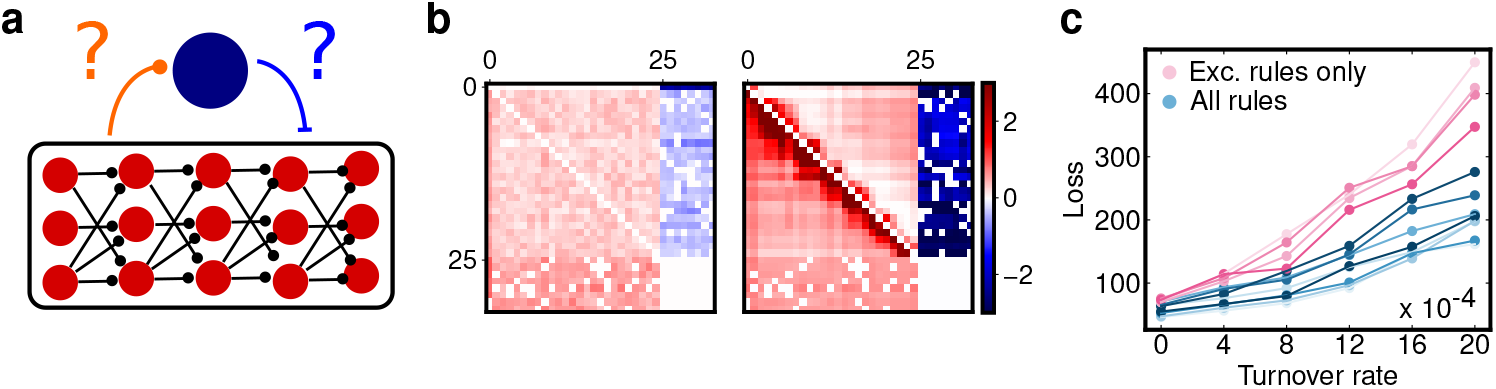
Learning plasticity on all synapses. **(a)** Schematic of synapses upon which we newly allow plasticity. **(b)** Weight matrices on activations 1 and 400 of a network evolving under plasticity rules learned on all 3 sets of synapses. I synapses increase. **(c)** Comparison of performance of rules learned only on E→E synapses (red) (N=5) versus all sets of synapses (blue) (N=8) when training includes synaptic turnover across varying rates of synaptic turnover.

#### 2.3.1 E→I and I→E synaptic plasticity improves decoding of time, particularly in the presence of synaptic turnover

Meta-learning uncovered triplets of plasticity rules that successfully organized initially random networks into sequence generators (Fig. 5b). When trained with turnover in E→E synapses, solutions that acted upon E→I and I→E synapses in addition to E→E outperformed solutions that only acted upon E→E connections, particularly when the rate of synaptic turnover was high (Fig. 5c), suggesting this additional plasticity played an important role in maintaining network dynamics through perturbation.

To investigate how rules acting on all synapses generated improved time representations, we repeated the dropout analysis. We found that E→E plasticity within these learned triples was largely similar to the rules previously learned on E→E synapses alone (Eqs. 4 and 6): solutions were consistently sensitive to the removal of 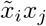 and 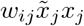,which appeared consistently with positive and negative coefficients, respectively. Further, dependence on these terms persisted when we trained networks with turnover on E→E synapses or I→E synapses (Supp. Fig. 6). As expected, we also found that solutions depended heavily upon terms that acted upon E→I and I→E synapses, but this plasticity was more difficult to interpret due to increased trial to trial variability in the discovered rules. We found, however, that the E→I plasticity rule consistently depended on the second order presynaptic term 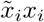,which always appeared with a positive coefficient, suggesting that E cells project to inhibitory counterparts with a strength that increases with the E cell level of activity. An implication of dependence on this term is that the strength of an E neuron’s recurrent inhibition, defined as 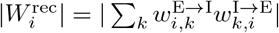, where *i* is the index of the E cell and *k* indexes the I cells to which it projects, might depend on its level of activity. Thus, ablation of excitatory inputs to an E cell might cause its recurrent, disynaptic inhibition to lower in a manner that homeostatically restores its firing.

**Figure 6:**
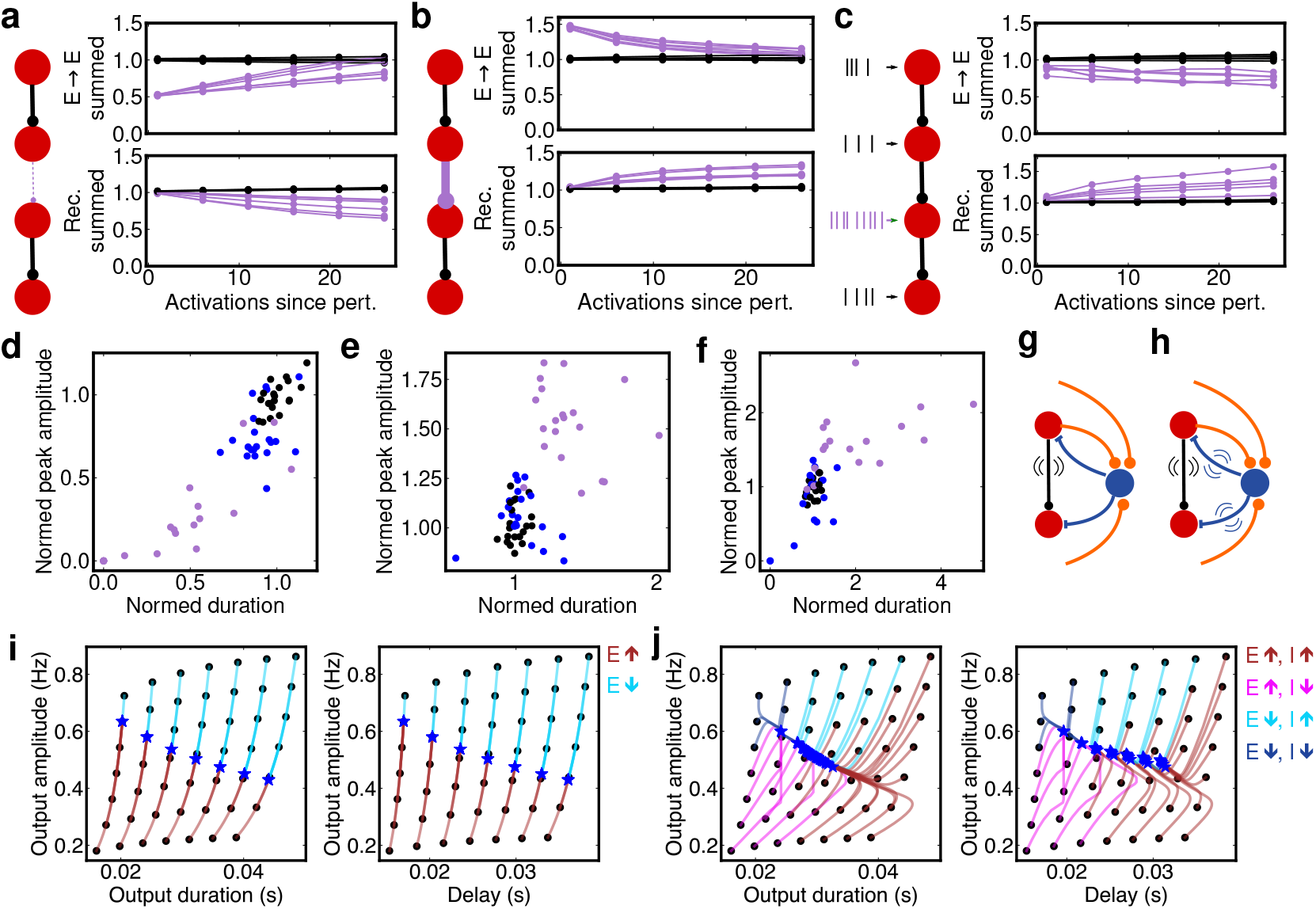
Network perturbations reveal homeostatic compensation in E→E and I→E synapses. **(a)** Response of the magnitude of summed E→E weights 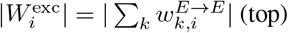 (top) and summed recurrent weight 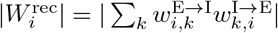 (bottom) to the imposed scaling down of E→ E weights to a single cell. Values for neuron with perturbed inputs shown in light purple; all others black. Each line represents average values for a single learned rule over N=20 networks. **(b)** Same as (a), but for imposed upscaling of E→ E weights to one cell (light purple). **(c)** Same as (a), but frequency of stochastic inputs to one cell is greatly increased (light purple) relative to all other cells (black). **(d-f)** Postsynaptic response of affected neurons in 20 networks plotted in the space of normalized peak amplitude and duration for manipulations (a-c). Black dots represent pre-perturbation responses; light purple, immediately after perturbation; blue, 25 activations after perturbation. **(g)** Schematic of feed-forward motif maintaining homeostasis via its E input. **(h)** Schematic of feed-forward motif with homeostasis on E→E and I→E synapses. **(i)** Phase flow in space of output firing envelope duration and amplitude (left) for a single neuron. Black points are first responses of the neuron for various inputs; blue stars represent responses after 200 activations. Note: initial and final output durations and delays are roughly the same. Colors of trajectories indicate how the neuron’s input weight changes to satisfy the plasticity rule: see legend. **(j)** If the forms of homeostasis on E→E and I→E maintain different aspects of the postsynaptic response, an attractor forms in the duration and peak amplitude of the postsynaptic response (left). The response delay is also constrained (right).

#### 2.3.2 Recurrent inhibition of E neurons is homeostatic in networks with learned plasticity

As the role of E→I and I→E plasticity was not completely clarified by our perturbations so far, we next investigated how this plasticity adjusted synapses coupled to an individual E cell when its typical input was manipulated. Noting the dependence of the E→I plasticity rule on 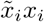,we hypothesized that a targeted neuron’s recurrent inhibition and excitatory afferents might be adjusted in concert to restore its typical firing pattern. We used discovered learning rules to organize sequences and then performed three varieties of *in silico* manipulations of individual E cells within these networks. In the first, we scaled down the excitatory afferents to the targeted E cell by 50% (Fig. 6a, diagram). In the second, we scaled up the same connections by 50% (Fig. 6b). In the last, we increased the rate of the targeted cell’s input Poisson process by a factor of 10 (Fig. 6c). This final manipulation mirrored the viral insertion of NaChBac to HVC_RA_ neurons in Wang et al., which causes these cells to become hyper-excitable [18, 45, 46]. Wang et al. found that manipulated cells recruited additional inhibition and weakened excitatory afferents.

We found that scaling down a targeted E cell’s excitatory afferents resulted in a rescaling of those excitatory weights and an accompanying decrease in 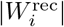 (Fig. 6a). We further found that these synaptic changes restore the initial firing pattern of the E cell: in Fig 6d, we plot the initial responses of targeted E cells across 20 self-organized sequences in the space of duration and peak amplitude of response (black points). Scaling down the excitatory afferents initially causes both the peak amplitude and duration of response to decrease (Fig. 6d; light purple points), but these responses largely recover after ∼25 activations of the network (Fig. 6d; blue points). When we instead strengthened E afferents to the targeted cell, we found these connections weakened and recurrent inhibition strengthened in compensatory fashion (Fig. 6b). In these networks, we found that targeted E cell responses that were initially lengthened with increased peak amplitude (Fig. 6e; light purple points) were reduced back to their pre-perturbation values of duration and amplitude (Fig. 6e; black points show initial responses; blue points, responses after 25 activations). Finally, rendering the targeted E cell hyper-excitable caused it to scale down its excitatory afferents and increase its recurrent inhibition in a manner that restored its typical firing pattern. In summary, plasticity rules on excitatory afferents and recurrent inhibition operate in tandem to maintain the firing pattern of the neuron (see Supp. Sec. 7 for additional details).

#### 2.3.3 Two forms of homeostasis create an attractor in postsynaptic response and timing

How might modulation of recurrent inhibition contribute differently to the modulation of excitation in preserving network dynamics? We hypothesized that distinct plasticity on different sets of synapses might confer a robust representation of time if these rules governed distinct aspects of the desired network function. For instance, within our HVC-like circuit, peak firing of E neurons might be controlled by E→E plasticity while total activity might be controlled by I→E plasticity. Since excitatory and inhibitory inputs to E cells differ in their timescales (E inputs are transient while I inputs are relatively tonic), we reasoned plasticity on both sets of synapses should increase the control of the postsynaptic response. While it is already known that multiple plasticity mechanisms can sharpen the responses of neurons to stimuli [37], prior work has not addressed whether similar plasticity might be leveraged to preserve the timing of such responses, which is crucial to timing representations. To explore this, we constructed a simplified model of a single neuron responding to a broad range of excitatory inputs, characterized by varying peak amplitudes and durations, and a tonic inhibitory input. We compared the responses of the neuron when both the excitatory and inhibitory input synapses (Fig. 6h) evolved under plasticity rules to responses produced when only the excitatory synapse evolved under a plasticity rule (Fig. 6g) and found the two rule model was able to better constrain the duration, peak amplitude, and timing of the postsynaptic response the E neuron (Fig. 6i-j; see Supp. Sec. 8 for full description of reduced model).

## 3 Discussion and limitations

Meta-learning plasticity rules via stochastic optimization is a promising technique, but suffers a number of limitations. One, the optimization process becomes expensive as the size of the rule basis, the number of neurons in the network, and amount of simulation time required grows. Further, CMA-ES may require many epochs to converge on good solutions. Training E→E plasticity (plasticity on all synapses) across 10 networks in batch typically required 24 (72) hours of compute on 30 Cascade Lake or Ice Lake Intel CPU cores to yield reasonable solutions. Two, meta-learning tends to generate different solutions based on the seed; due to the expensive nature of each trial, we did not carry out enough trials to claim full knowledge of the solution space. Three, the plasticity rules learned were quite dense in our choice of basis, limiting interpretation, and we ultimately employed perturbations to better understand the critical terms. Four, the choice of basis limits the space of discoverable rules; for instance, we did not include feedback-modulated plasticity in this study. Five, the initial connectivity of the circuit likely has a strong bearing on the sort of plasticity that successfully can leverage it [47, 48].

Prior to any experience, intrinsic, self-organized dynamics within the brain can serve as powerful priors that can accelerate and shape successful, feedback driven learning. In this work, we study how one such computational primitive could emerge by adapting a meta-learning procedure to learn the learning rules that self-organize and maintain robust representations of time in neural dynamics. Meta-learning discovers a temporally asymmetric (STDP-like) generalization of Oja’s rule that organizes and maintains sparse, sequential activity out of initially random connectivity, which outperforms other models of sequence generation in the presence of synaptic turnover by permitting flexible rescaling of inputs to restore dynamics. Additionally we found that plasticity rules learned on all sets of synapses outperform plasticity rules applied only to E connections. Through a toy model, we show how plasticity on all synapses could confer extra timing stability if the plasticity in distinct sets of synapses act on different moments of an E neuron’s activity.

In this work, we have developed a paradigm to understand how computational primitives might self-organize within neural circuits and selected sequences as our test example. Future work could study the emergence of other canonical forms of neural dynamics that have been widely identified in brain activity and serve as fundamental components of computation, such as line attractors or limit cycles, or how plasticity rules generating such components interact with rules requiring feedback.

## 4 Acknowledgements and disclosure of funding

We would like to thank Carlos Lois, Zsofia Török, Bo Wang, Patrick Zhang, and Leenoy Meshulam for useful discussions. This work was supported by the Simons Collaboration for the Global Brain and an NIH BRAIN grant (5R01NS104925). This work was facilitated through the use of advanced computational, storage, and networking infrastructure provided by the Hyak supercomputer system and funded by the STF at the University of Washington.

## NeurIPS Paper Checklist

### 1. Claims

Question: Do the main claims made in the abstract and introduction accurately reflect the paper’s contributions and scope?

Answer: [Yes]

Justification: The primary claims of the paper are addressed computationally in Section 2, results, and are expanded upon in the supplement.

Guidelines:

- The answer NA means that the abstract and introduction do not include the claims made in the paper.
- The abstract and/or introduction should clearly state the claims made, including the contributions made in the paper and important assumptions and limitations. A No or NA answer to this question will not be perceived well by the reviewers.
- The claims made should match theoretical and experimental results, and reflect how much the results can be expected to generalize to other settings.
- It is fine to include aspirational goals as motivation as long as it is clear that these goals are not attained by the paper.

### 2. Limitations

Question: Does the paper discuss the limitations of the work performed by the authors? Answer: [Yes]

Justification: The limitations of the findings and methods are discussed extensively in the Discussion and Limitations section.

Guidelines:

- The answer NA means that the paper has no limitation while the answer No means that the paper has limitations, but those are not discussed in the paper.
- The authors are encouraged to create a separate “Limitations” section in their paper.
- The paper should point out any strong assumptions and how robust the results are to violations of these assumptions (e.g., independence assumptions, noiseless settings, model well-specification, asymptotic approximations only holding locally). The authors should reflect on how these assumptions might be violated in practice and what the implications would be.
- The authors should reflect on the scope of the claims made, e.g., if the approach was only tested on a few datasets or with a few runs. In general, empirical results often depend on implicit assumptions, which should be articulated.
- The authors should reflect on the factors that influence the performance of the approach. For example, a facial recognition algorithm may perform poorly when image resolution is low or images are taken in low lighting. Or a speech-to-text system might not be used reliably to provide closed captions for online lectures because it fails to handle technical jargon.
- The authors should discuss the computational efficiency of the proposed algorithms and how they scale with dataset size.
- If applicable, the authors should discuss possible limitations of their approach to address problems of privacy and fairness.
- While the authors might fear that complete honesty about limitations might be used by reviewers as grounds for rejection, a worse outcome might be that reviewers discover limitations that aren’t acknowledged in the paper. The authors should use their best judgment and recognize that individual actions in favor of transparency play an important role in developing norms that preserve the integrity of the community. Reviewers will be specifically instructed to not penalize honesty concerning limitations.

### 3. Theory Assumptions and Proofs

Question: For each theoretical result, does the paper provide the full set of assumptions and a complete (and correct) proof?

Answer: [NA]

Justification: The paper sketches several theoretical ideas, but does not make rigorous theoretical claims, instead opting to illustrate phenomena via computational approaches.

Guidelines:

- The answer NA means that the paper does not include theoretical results.
- All the theorems, formulas, and proofs in the paper should be numbered and cross-referenced.
- All assumptions should be clearly stated or referenced in the statement of any theorems.
- The proofs can either appear in the main paper or the supplemental material, but if they appear in the supplemental material, the authors are encouraged to provide a short proof sketch to provide intuition.
- Inversely, any informal proof provided in the core of the paper should be complemented by formal proofs provided in appendix or supplemental material.
- Theorems and Lemmas that the proof relies upon should be properly referenced.

### 4. Experimental Result Reproducibility

Question: Does the paper fully disclose all the information needed to reproduce the main experimental results of the paper to the extent that it affects the main claims and/or conclusions of the paper (regardless of whether the code and data are provided or not)?

Answer: [Yes]

Justification: Code referenced in supplementary section, Code Availability.

- The answer NA means that the paper does not include experiments.
- If the paper includes experiments, a No answer to this question will not be perceived well by the reviewers: Making the paper reproducible is important, regardless of whether the code and data are provided or not.
- If the contribution is a dataset and/or model, the authors should describe the steps taken to make their results reproducible or verifiable.
- Depending on the contribution, reproducibility can be accomplished in various ways. For example, if the contribution is a novel architecture, describing the architecture fully might suffice, or if the contribution is a specific model and empirical evaluation, it may be necessary to either make it possible for others to replicate the model with the same dataset, or provide access to the model. In general. releasing code and data is often one good way to accomplish this, but reproducibility can also be provided via detailed instructions for how to replicate the results, access to a hosted model (e.g., in the case of a large language model), releasing of a model checkpoint, or other means that are appropriate to the research performed.
- While NeurIPS does not require releasing code, the conference does require all submissions to provide some reasonable avenue for reproducibility, which may depend on the nature of the contribution. For example
  a. If the contribution is primarily a new algorithm, the paper should make it clear how to reproduce that algorithm.
  b. If the contribution is primarily a new model architecture, the paper should describe the architecture clearly and fully.
  c. If the contribution is a new model (e.g., a large language model), then there should either be a way to access this model for reproducing the results or a way to reproduce the model (e.g., with an open-source dataset or instructions for how to construct the dataset).
  d. We recognize that reproducibility may be tricky in some cases, in which case authors are welcome to describe the particular way they provide for reproducibility. In the case of closed-source models, it may be that access to the model is limited in some way (e.g., to registered users), but it should be possible for other researchers to have some path to reproducing or verifying the results.

### 5. Open access to data and code

Question: Does the paper provide open access to the data and code, with sufficient instructions to faithfully reproduce the main experimental results, as described in supplemental material?

Answer: [Yes]

Justification: Code referenced in supplementary section, Code Availability. Guidelines:

- The answer NA means that paper does not include experiments requiring code.
- Please see the NeurIPS code and data submission guidelines (https://nips.cc/public/guides/CodeSubmissionPolicy) for more details.
- While we encourage the release of code and data, we understand that this might not be possible, so “No” is an acceptable answer. Papers cannot be rejected simply for not including code, unless this is central to the contribution (e.g., for a new open-source benchmark).
- The instructions should contain the exact command and environment needed to run to reproduce the results. See the NeurIPS code and data submission guidelines (https://nips.cc/public/guides/CodeSubmissionPolicy) for more details.
- The authors should provide instructions on data access and preparation, including how to access the raw data, preprocessed data, intermediate data, and generated data, etc.
- The authors should provide scripts to reproduce all experimental results for the new proposed method and baselines. If only a subset of experiments are reproducible, they should state which ones are omitted from the script and why.
- At submission time, to preserve anonymity, the authors should release anonymized versions (if applicable).
- Providing as much information as possible in supplemental material (appended to the paper) is recommended, but including URLs to data and code is permitted.

### 6. Experimental Setting/Details

Question: Does the paper specify all the training and test details (e.g., data splits, hyperparameters, how they were chosen, type of optimizer, etc.) necessary to understand the results?

Answer: [Yes]

Justification: The paper clearly outlines all methods. See section: Results, and also supplement.

Guidelines:

- The answer NA means that the paper does not include experiments.
- The experimental setting should be presented in the core of the paper to a level of detail that is necessary to appreciate the results and make sense of them.
- The full details can be provided either with the code, in appendix, or as supplemental material.

### 7. Experiment Statistical Significance

Question: Does the paper report error bars suitably and correctly defined or other appropriate information about the statistical significance of the experiments?

Answer: [Yes]

Justification: Section 2, results. Guidelines:

- The answer NA means that the paper does not include experiments.
- The authors should answer “Yes” if the results are accompanied by error bars, confidence intervals, or statistical significance tests, at least for the experiments that support the main claims of the paper.
- The factors of variability that the error bars are capturing should be clearly stated (for example, train/test split, initialization, random drawing of some parameter, or overall run with given experimental conditions).
- The method for calculating the error bars should be explained (closed form formula, call to a library function, bootstrap, etc.)
- The assumptions made should be given (e.g., Normally distributed errors).
- It should be clear whether the error bar is the standard deviation or the standard error of the mean.
- It is OK to report 1-sigma error bars, but one should state it. The authors should preferably report a 2-sigma error bar than state that they have a 96% CI, if the hypothesis of Normality of errors is not verified.
- For asymmetric distributions, the authors should be careful not to show in tables or figures symmetric error bars that would yield results that are out of range (e.g. negative error rates).
- If error bars are reported in tables or plots, The authors should explain in the text how they were calculated and reference the corresponding figures or tables in the text.

### 8. Experiments Compute Resources

Question: For each experiment, does the paper provide sufficient information on the computer resources (type of compute workers, memory, time of execution) needed to reproduce the experiments?

Answer: [Yes]

Justification: The computational resources used are detailed in the Discussion section. Guidelines:

- The answer NA means that the paper does not include experiments.
- The paper should indicate the type of compute workers CPU or GPU, internal cluster, or cloud provider, including relevant memory and storage.
- The paper should provide the amount of compute required for each of the individual experimental runs as well as estimate the total compute.
- The paper should disclose whether the full research project required more compute than the experiments reported in the paper (e.g., preliminary or failed experiments that didn’t make it into the paper).

### 9. Code Of Ethics

Question: Does the research conducted in the paper conform, in every respect, with the NeurIPS Code of Ethics https://neurips.cc/public/EthicsGuidelines?

Answer: [Yes]

Justification: The NeurIPS code of ethics is fully abided by in this work. Guidelines:

- The answer NA means that the authors have not reviewed the NeurIPS Code of Ethics.
- If the authors answer No, they should explain the special circumstances that require a deviation from the Code of Ethics.
- The authors should make sure to preserve anonymity (e.g., if there is a special consideration due to laws or regulations in their jurisdiction).

### 10. Broader Impacts

Question: Does the paper discuss both potential positive societal impacts and negative societal impacts of the work performed?

Answer: [Yes]

Justification: Societal impacts are expounded upon in the discussion section.

Guidelines:

- The answer NA means that there is no societal impact of the work performed.
- If the authors answer NA or No, they should explain why their work has no societal impact or why the paper does not address societal impact.
- Examples of negative societal impacts include potential malicious or unintended uses (e.g., disinformation, generating fake profiles, surveillance), fairness considerations (e.g., deployment of technologies that could make decisions that unfairly impact specific groups), privacy considerations, and security considerations.
- The conference expects that many papers will be foundational research and not tied to particular applications, let alone deployments. However, if there is a direct path to any negative applications, the authors should point it out. For example, it is legitimate to point out that an improvement in the quality of generative models could be used to generate deepfakes for disinformation. On the other hand, it is not needed to point out that a generic algorithm for optimizing neural networks could enable people to train models that generate Deepfakes faster.
- The authors should consider possible harms that could arise when the technology is being used as intended and functioning correctly, harms that could arise when the technology is being used as intended but gives incorrect results, and harms following from (intentional or unintentional) misuse of the technology.
- If there are negative societal impacts, the authors could also discuss possible mitigation strategies (e.g., gated release of models, providing defenses in addition to attacks, mechanisms for monitoring misuse, mechanisms to monitor how a system learns from feedback over time, improving the efficiency and accessibility of ML).

### 11. Safeguards

Question: Does the paper describe safeguards that have been put in place for responsible release of data or models that have a high risk for misuse (e.g., pretrained language models, image generators, or scraped datasets)?

Answer: [NA]

Justification: No risk of misuse.

Guidelines:

- The answer NA means that the paper poses no such risks.
- Released models that have a high risk for misuse or dual-use should be released with necessary safeguards to allow for controlled use of the model, for example by requiring that users adhere to usage guidelines or restrictions to access the model or implementing safety filters.
- Datasets that have been scraped from the Internet could pose safety risks. The authors should describe how they avoided releasing unsafe images.
- We recognize that providing effective safeguards is challenging, and many papers do not require this, but we encourage authors to take this into account and make a best faith effort.

### 12. Licenses for existing assets

Question: Are the creators or original owners of assets (e.g., code, data, models), used in the paper, properly credited and are the license and terms of use explicitly mentioned and properly respected?

Answer: [Yes]

Justification: Code, data, and models used in the study belong to the authors or are properly credited.

Guidelines:

- The answer NA means that the paper does not use existing assets.
- The authors should cite the original paper that produced the code package or dataset.
- The authors should state which version of the asset is used and, if possible, include a URL.
- The name of the license (e.g., CC-BY 4.0) should be included for each asset.
- For scraped data from a particular source (e.g., website), the copyright and terms of service of that source should be provided.
- If assets are released, the license, copyright information, and terms of use in the package should be provided. For popular datasets, paperswithcode.com/datasets has curated licenses for some datasets. Their licensing guide can help determine the license of a dataset.
- For existing datasets that are re-packaged, both the original license and the license of the derived asset (if it has changed) should be provided.
- If this information is not available online, the authors are encouraged to reach out to the asset’s creators.

### 13. New Assets

Question: Are new assets introduced in the paper well documented and is the documentation provided alongside the assets?

Answer: [Yes]

Justification: All new assets are well documented.

Guidelines:

- The answer NA means that the paper does not release new assets.
- Researchers should communicate the details of the dataset/code/model as part of their submissions via structured templates. This includes details about training, license, limitations, etc.
- The paper should discuss whether and how consent was obtained from people whose asset is used.
- At submission time, remember to anonymize your assets (if applicable). You can either create an anonymized URL or include an anonymized zip file.

### 14. Crowdsourcing and Research with Human Subjects

Question: For crowdsourcing experiments and research with human subjects, does the paper include the full text of instructions given to participants and screenshots, if applicable, as well as details about compensation (if any)?

Answer: [NA]

Justification: No human subjected were used in this study.

Guidelines:

- The answer NA means that the paper does not involve crowdsourcing nor research with human subjects.
- Including this information in the supplemental material is fine, but if the main contribution of the paper involves human subjects, then as much detail as possible should be included in the main paper.
- According to the NeurIPS Code of Ethics, workers involved in data collection, curation, or other labor should be paid at least the minimum wage in the country of the data collector.

### 15. Institutional Review Board (IRB) Approvals or Equivalent for Research with Human Subjects

Question: Does the paper describe potential risks incurred by study participants, whether such risks were disclosed to the subjects, and whether Institutional Review Board (IRB) approvals (or an equivalent approval/review based on the requirements of your country or institution) were obtained?

Answer: [NA]

Justification: [NA]

Guidelines: No IRB approvals were required for this study.

- The answer NA means that the paper does not involve crowdsourcing nor research with human subjects.
- Depending on the country in which research is conducted, IRB approval (or equivalent) may be required for any human subjects research. If you obtained IRB approval, you should clearly state this in the paper.
- We recognize that the procedures for this may vary significantly between institutions and locations, and we expect authors to adhere to the NeurIPS Code of Ethics and the guidelines for their institution.
- For initial submissions, do not include any information that would break anonymity (if applicable), such as the institution conducting the review.

## References

[1] W Nicola and C Clopath. Supervised learning in spiking neural networks with force training. Nat Commun, 8(1):2208, 2017. ISSN 2041-1723. doi: 10.1038/s41467-017-01827-3.

[2] Elias Najarro and Sebastian Risi. Meta-Learning through Hebbian Plasticity in Random Networks, April 2022. URL http://arxiv.org/abs/2007.02686. 2007.02686.

[3] Nikhil X Bhattasali, Anthony M Zador, and Tatiana A Engel. Neural Circuit Architectural Priors for Embodied Control.

[4] Nikhil X. Bhattasali, Venkatesh Pattabiraman, Lerrel Pinto, and Grace W. Lindsay. Neural Circuit Architectural Priors for Quadruped Locomotion, October 2024. URL http://arxiv.org/abs/2410.07174. 2410.07174.

[5] Jane X Wang, Zeb Kurth-Nelson, Dhruva Tirumala, Hubert Soyer, Joel Z Leibo, Remi Munos, Charles Blundell, Dharshan Kumaran, and Matt Botvinick. Learning to reinforcement learn. arXiv preprint arXiv:1611.05763, 2016.

[6] Pastalkova E, Itskov V, Amarasingham A, and Buzsáki G. Internally generated cell assembly sequences in the rat hippocampus. Science, 321(5894):1322–7, sep 2008. doi: 10.1126/science.1159775.

[7] Luczak A, Barthó P, Marguet SL, Buzsáki G, and Harris KD. Sequential structure of neocortical spontaneous activity in vivo. Proc Natl Acad Sci, 104(1), 2007. doi: 10.1073/pnas.0605643104.

[8] Gustavo B.M Mello, Sofia Soares, and Joseph J Paton. A Scalable Population Code for Time in the Striatum. Current Biology, 25(9):1113–1122, may 2015. ISSN 0960-9822. doi: 10.1016/j.cub.2015.02.036. URL https://doi.org/10.1016/j.cub.2015.02.036. Publisher: Elsevier.

[9] R. H. Hahnloser, A. A. Kozhevnikov, and M. S. Fee. An ultra-sparse code underlies the generation of neural sequences in a songbird. Nature, 419(6902):65–70, 2002. ISSN 0028-0836 (Print) 0028-0836. doi: 10.1038/nature00974.

[10] Ila R. Fiete, Richard H.R. Hahnloser, Michale S. Fee, and H. Sebastian Seung. Temporal sparseness of the premotor drive is important for rapid learning in a neural network model of birdsong. Journal of neurophysiology, 92:2274–2282, 2004. doi: 10.1152/jn.01133.2003.

[11] IR Fiete, W Senn, CZ Wang, and RH Hahnloser. Spike-time-dependent plasticity and heterosy-naptic competition organize networks to produce long scale-free sequences of neural activity. Neuron, 65:563–76, 2010. doi: 10.1016/j.neuron.2010.02.003.

[12] J Jun and D Jin. Development of Neural Circuitry for Precise Temporal Sequences through Spontaneous Activity, Axon Remodeling, and Synaptic Plasticity. PLOS ONE, 2(8):e723, August 2007. ISSN 1932-6203. doi: 10.1371/journal.pone.0000723. URL https://journals.plos.org/plosone/article?id=10.1371/journal.pone.0000723. Publisher: Public Library of Science.

[13] P. Zheng and J. Triesch. Robust development of synfire chains from multiple plasticity mechanisms. Front Comput Neurosci, 8:66, 2014. ISSN 1662-5188 (Print) 1662-5188. doi: 10.3389/fncom.2014.00066.

[14] Murray JM and Escola GS. Learning multiple variable-speed sequences in striatum via cortical tutoring. Elife, 6, 2017. doi: 10.7554/eLife.26084.

[15] Felix Weissenberger, Florian Meier, Johannes Lengler, Hafsteinn Einarsson, and Angelika Steger. Long synfire chains emerge by spike-timing dependent plasticity modulated by population activity. International Journal of Neural Systems, 27(08):1750044, 2017. doi: 10.1142/S0129065717500447. URL https://doi.org/10.1142/S0129065717500447. PMID: 28982282.

[16] U. Pereira and N. Brunel. Unsupervised learning of persistent and sequential activity. Front Comput Neurosci, 13:97, 2019. ISSN 1662-5188 (Print) 1662-5188. doi: 10.3389/fncom.2019.00097.

[17] Yevhen Tupikov and Dezhe Z Jin. Addition of new neurons and the emergence of a local neural circuit for precise timing. page 43, 2021.

[18] B. Wang, Z. Torok, A. Duffy, D. G. Bell, S. Wongso, T. A. F. Velho, A. L. Fairhall, and C. Lois. Unsupervised restoration of a complex learned behavior after large-scale neuronal perturbation. Nat Neurosci, 2024. ISSN 1097-6256. doi: 10.1038/s41593-024-01630-6.

[19] Y. Bengio, S. Bengio, and J. Cloutier. Learning a synaptic learning rule. ii:969 vol.2–, 1991. doi: 10.1109/IJCNN.1991.155621.

[20] N. Shervani-Tabar and R. Rosenbaum. Meta-learning biologically plausible plasticity rules with random feedback pathways. Nat Commun, 14(1):1805, 2023. ISSN 2041-1723. doi: 10.1038/s41467-023-37562-1.

[21] D. Tyulmankov, G. R. Yang, and L. F. Abbott. Meta-learning synaptic plasticity and memory addressing for continual familiarity detection. Neuron, 110(3):544–557.e8, 2022. ISSN 0896-6273 (Print) 0896-6273. doi: 10.1016/j.neuron.2021.11.009.

[22] Samy Bengio, Yoshua Bengio, Jocelyn Cloutier, and Jan Gecsei. On the optimization of a synaptic learning rule. In Optimality in Biological and Artificial Networks?, pages 265–287. Routledge, 2013.

[23] Jack Lindsey and Ashok Litwin-Kumar. Learning to Learn with Feedback and Local Plasticity. In H. Larochelle, M. Ranzato, R. Hadsell, M. F. Balcan, and H. Lin, editors, Advances in Neural Information Processing Systems, volume 33, pages 21213–21223. Curran Associates, Inc., 2020. URL https://proceedings.neurips.cc/paper_files/paper/2020/file/f291e10ec3263bd7724556d62e70e25d-Paper.pdf.

[24] Luke Metz, Niru Maheswaranathan, Brian Cheung, and Jascha Sohl-Dickstein. Meta-Learning Update Rules for Unsupervised Representation Learning, February 2019. URL http://arxiv.org/abs/1804.00222. 1804.00222.

[25] Thomas Miconi, Jeff Clune, and Kenneth O. Stanley. Differentiable plasticity: training plastic neural networks with backpropagation, July 2018. URL http://arxiv.org/abs/1804.02464. 1804.02464.

[26] Hector Garcia Rodriguez, Qinghai Guo, and Timoleon Moraitis. Short-Term Plasticity Neurons Learning to Learn and Forget, June 2022. URL http://arxiv.org/abs/2206.14048. 2206.14048.

[27] Michael A. Long, Dezhe Z. Jin, and Michale S. Fee. Support for a synaptic chain model of neuronal sequence generation. Nature, 468(7322):394–399, November 2010. ISSN 0028-0836, 1476-4687. doi: 10.1038/nature09514. URL http://www.nature.com/articles/nature09514.

[28] F. W. Moll, D. Kranz, A. Corredera Asensio, et al. Thalamus drives vocal onsets in the zebra finch courtship song. Nature, 616:132–136, 2023. doi: 10.1038/s41586-023-05818-x. URL https://doi.org/10.1038/s41586-023-05818-x.

[29] H. H. Danish, D. Aronov, and M. S. Fee. Rhythmic syllable-related activity in a songbird motor thalamic nucleus necessary for learned vocalizations. PLoS ONE, 12(6):e0169568, 2017. doi: 10.1371/journal.pone.0169568.

[30] G. Kosche, D. Vallentin, and M. A. Long. Interplay of inhibition and excitation shapes a premotor neural sequence. J Neurosci, 35(3):1217–27, 2015. ISSN 0270-6474 (Print) 0270-6474. doi: 10.1523/jneurosci.4346-14.2015.

[31] Barish Poole, Jeffrey E. Markowitz, and Timothy J. Gardner. The Song Must Go On: Resilience of the Songbird Vocal Motor Pathway. PLoS ONE, 7(6):e38173, June 2012. doi: 10.1371/journal.pone.0038173. URL https://pmc.ncbi.nlm.nih.gov/articles/PMC3387175/.

[32] Richard Mooney and Jonathan F. Prather. The HVC Microcircuit: The Synaptic Basis for Interactions between Song Motor and Vocal Plasticity Pathways. Journal of Neuroscience, 25(8):1952–1964, 2005. ISSN 0270-6474. doi: 10.1523/JNEUROSCI.3726-04.2005. URL https://www.jneurosci.org/content/25/8/1952. Publisher: Society for Neuroscience _eprint: https://www.jneurosci.org/content/25/8/1952.full.pdf.

[33] Basile Confavreux, Friedemann Zenke, Everton J. Agnes, Timothy Lillicrap, and Tim P. Vogels. A meta-learning approach to (re)discover plasticity rules that carve a desired function into a neural network. In 34th Conference on Neural Information Processing Systems, Vancouver, Canada, 2020.

[34] Anne Auger and Nikolaus Hansen. A restart cma evolution strategy with increasing population size. volume 2, pages 1769–1776, 01 2005. doi: 10.1109/CEC.2005.1554902.

[35] Manuel A. Castro-Alamancos and Barry W. Connors. Distinct forms of short-term plasticity at excitatory synapses of hippocampus and neocortex. Proceedings of the National Academy of Sciences, 94(8):4161–4166, April 1997. doi: 10.1073/pnas.94.8.4161. URL https://www.pnas.org/doi/10.1073/pnas.94.8.4161. Publisher: Proceedings of the National Academy of Sciences.

[36] Rylan S. Larsen and P. Jesper Sjöström. Synapse-type-specific plasticity in local circuits. Current opinion in neurobiology, 35:127, August 2015. doi: 10.1016/j.conb.2015.08.001. URL https://pmc.ncbi.nlm.nih.gov/articles/PMC5280068/.

[37] T. P. Vogels, H. Sprekeler, F. Zenke, C. Clopath, and W. Gerstner. Inhibitory plasticity balances excitation and inhibition in sensory pathways and memory networks. Science, 334(6062):1569–73, 2011. ISSN 0036-8075. doi: 10.1126/science.1211095.

[38] Ashok Litwin-Kumar and Brent Doiron. Formation and maintenance of neuronal assemblies through synaptic plasticity. Nature communications, 5(1):5319, 2014.

[39] Yotam Luz and Maoz Shamir. Balancing Feed-Forward Excitation and Inhibition via Hebbian Inhibitory Synaptic Plasticity. PLOS Computational Biology, 8(1):1–12, January 2012. doi: 10.1371/journal.pcbi.1002334. URL https://doi.org/10.1371/journal.pcbi.1002334. Publisher: Public Library of Science.

[40] Waitzmann F, Wu YK, and Gjorgjieva J. Top-down modulation in canonical cortical circuits with short-term plasticity. Proc Natl Acad Sci, 121(16), 2024. doi: 10.1073/pnas.2311040121.

[41] Ziyi Gong and Nicolas Brunel. Inhibitory Plasticity Enhances Sequence Storage Capacity and Retrieval Robustness, April 2024. URL https://www.biorxiv.org/content/10.1101/2024.04.08.588573v1. Pages: 2024.04.08.588573 Section: New Results.

[42] Fereshteh Lagzi and Adrienne L. Fairhall. Emergence of co-tuning in inhibitory neurons as a network phenomenon mediated by randomness, correlations, and homeostatic plasticity. Science Advances, 10(12):eadi4350, March 2024. ISSN 2375-2548. doi: 10.1126/sciadv.adi4350. URL https://www.science.org/doi/10.1126/sciadv.adi4350.

[43] Basile Confavreux, Everton J. Agnes, Friedemann Zenke, Henning Sprekeler, and Tim P. Vogels. Balancing complexity, performance and plausibility to meta learn plasticity rules in recurrent spiking networks, June 2024. URL https://www.biorxiv.org/content/10.1101/2024.06.17.599260v1. Pages: 2024.06.17.599260 Section: New Results.

[44] Basile Confavreux, Poornima Ramesh, Pedro J Gonçalves, Jakob H Macke, and Tim P Vogels. Meta-learning families of plasticity rules in recurrent spiking networks using simulation-based inference.

[45] Shuyin Sim, Salome Antolin, Chia-Wei Lin, Yingxi Lin, and Carlos Lois. Increased cell-intrinsic excitability induces synaptic changes in new neurons in the adult dentate gyrus that require npas4. Journal of Neuroscience, 33(18):7928–7940, 2013. ISSN 0270-6474. doi: 10.1523/JNEUROSCI.1571-12.2013. URL https://www.jneurosci.org/content/33/18/7928.

[46] C. W. Lin, S. Sim, A. Ainsworth, M. Okada, W. Kelsch, and C. Lois. Genetically increased cell-intrinsic excitability enhances neuronal integration into adult brain circuits. Neuron, 65(1): 32–39, jan 2010. doi: 10.1016/j.neuron.2009.12.001.

[47] Ashok Litwin-Kumar, Kameron Decker Harris, Richard Axel, Haim Sompolinsky, and L. F. Abbott. Optimal Degrees of Synaptic Connectivity. Neuron, 93(5):1153–1164.e7, March 2017. ISSN 1097-4199. doi: 10.1016/j.neuron.2017.01.030.

[48] Kaushik J. Lakshminarasimhan, Marjorie Xie, Jeremy D. Cohen, Britton A. Sauerbrei, Adam W. Hantman, Ashok Litwin-Kumar, and Sean Escola. Specific connectivity optimizes learning in thalamocortical loops. Cell Reports, 43(4):114059, April 2024. ISSN 2211-1247. doi: 10.1016/j.celrep.2024.114059.

