## supplemental_info for "Discovering plasticity rules that organize and maintain neural circuits"

#### 1 Network simulations

##### 1.1 Network initialization

For all network simulations, network were initialized with excitatory units connected all-to-all with weights drawn from $U[0.2\,w^{(0)}_{E\to E}, w^{(0)}_{E\to E}]$. E→I and I→E weights were nonzero with probability 0.8 with values drawn from $\mathcal{N}(w^{(0)}_{E\to I}, 0.3\,w^{(0)}_{E\to I})$ and $\mathcal{N}(w^{(0)}_{I\to E}, 0.3\,|w^{(0)}_{I\to E}|)$, respectively. Any weights violating Dale's law were set to zero.

##### 1.2 Single neuron dynamics

All neurons were modeled as rate-based units with threshold linear activations, with continuous firing rates evolving according to

$$x_j(t) = [V_j(t) - b]^+\,, \tag{1}$$

where $V_j(t)$ evolved via

$$\tau_m \dot{V}_j(t) = -V_j(t) + \sum_i w_{ij} x_i(t) \tag{2}$$

Here, $w_{ij}$ is the weight of the synapse $i \to j$, $\tau_m$ is the membrane time constant, and $b$ is the bias. Values of $\tau_m$ differed across cell types (E and I), but were consistent within cell type. Eqs. 1 and 2 were solved numerically with timestep size $\delta t = 0.1$ ms.

##### 1.3 Network input

We adopt the assumptions made in Jun and Jin [5] and Tupikov and Jin [7] and drive a single neuron with a strong kick of excitation at the start of each trial. Following this, all other neurons in the network received two filtered Poisson process inputs, a "frozen" (stereotyped activation to activation) input and a stochastic input. These processes had rates $\lambda_{\text{frozen}} = \lambda^{(0)} p_{\text{fix}}$ and $\lambda_{\text{stoch}} = \lambda^{(0)}(1 - p_{\text{fix}})$, respectively, where $p_{\text{fix}} = 0.75$ and $\lambda^{(0)} = 80$ Hz. Arrivals from both processes were then smoothed into an input current via an alpha function kernel such that arrivals $\{t_i\}$ became input current

$$I(t) = \sum_{\{t_i\}} ae \frac{t - t_i}{\tau_{\text{input}}} e^{-(t-t_i)/\tau_{\text{input}}} \Theta(t - t_i), \tag{3}$$

where $\tau_{\text{input}} = 3$ ms and $a$ was 0.09 and 0.02 for excitatory and inhibitory neurons, respectively. Input current to the initially driven neuron was modeled in a similar fashion: a block of 10 contiguous arrivals was transformed into a current via Eq. 3.

38th Conference on Neural Information Processing Systems (NeurIPS 2024).

Table 1: Environment parameters

| Parameter | Description | Value |
|---|---|---|
| $w^{(0)}_{E\to E}$ | Maximum initial E→E weight | 0.4 |
| $w^{(0)}_{E\to I}$ | Mean initial E→I weight | 0.5 |
| $w^{(0)}_{I\to E}$ | Mean initial I→E weight | -0.3 |
| $\tau_m^{(E)}$ | Excitatory cell membrane time constant | 10 ms |
| $\tau_m^{(I)}$ | Inhibitory cell membrane time constant | 0.1 ms |
| $b$ | Excitatory cell resting potential | 0.1 |

#### 2 Meta-learning basis

To discover plasticity rules, we leverage a meta-learning approach pioneered by Bengio et al.[2] and more recently extended by Confavreux et al[3]. Specifically, we parameterize a set of plasticity rules with coefficients $c_k$ and time constants $\tau_k$, such that individual synapses evolve according to

$$\dot{w}_i = \sum_k c_k F_k(x_i, y_i, w_i, \tau_k) \tag{4}$$

where $x_i$ and $y_i$ are the pre and postsynaptic activity of synapse $i$, respectively, $w_i$ is the synapse size, and $F_k$ is the $k^{th}$ term in the plasticity rule. The terms of $\mathbf{F}$ are

$$\mathbf{F}(x_i, y_i, w_i) = \left\{ \begin{array}{c} 1 \\ y_i \\ x_i \\ x_i\, y_i \\ \tilde{y}_i^{(4)} \\ x_i\, \tilde{y}_i^{(5)} \\ \tilde{x}_i^{(6)} \\ \tilde{x}_i^{(7)}\, y_i \\ \tilde{y}_i^{(8)}\, y_i \\ \tilde{x}_i^{(9)}\, x_i \\ w_i \\ w_i y_i \\ w_i x_i \\ w_i x_i\, y_i \\ w_i \tilde{y}_i^{(14)} \\ w_i x_i\, \tilde{y}_i^{(15)} \\ w_i \tilde{x}_i^{(16)} \\ w_i \tilde{x}_i^{(17)}\, y_i \\ w_i \tilde{y}_i^{(18)}\, y_i \\ w_i \tilde{x}_i^{(19)}\, x_i \end{array} \right\}, \tag{5}$$

where

$$\tilde{x}_i^{(k)}(t) = \int_0^t e^{-(t-t')/\tau_k} x_i(t')dt', \tag{6}$$

$$\tilde{y}_i^{(k)}(t) = \int_0^t e^{-(t-t')/\tau_k} y_i(t')dt', \tag{7}$$

and $\tau_k$ is a learned time constant specific to element $k$ of $\mathbf{F}$. Note, we rescaled terms that were higher order in pre- or postsynaptic activity by a factor 0.1 per order in order to limit the number exploding simulations.

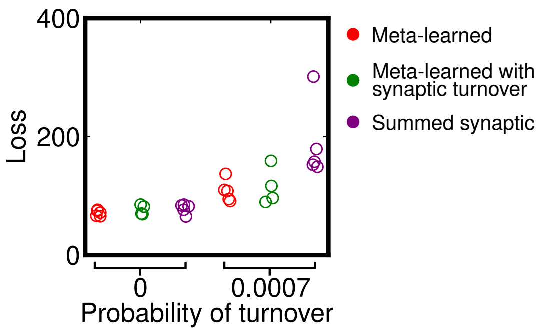


Figure 1: Five best versions of meta-learned rules trained with and without synaptic turnover compared to summed synaptic learning rule. Points represent median loss for each condition.

#### 3 Optimization via CMA-ES

To optimize rule coefficients and time constants, we used Covariance Matrix Adaptation (CMA-ES) [1], a technique that iteratively samples from a high dimensional Gaussian placed on the space of plasticity rule parameters. The mean of the Gaussian was initialized to 0 for all rule coefficient parameters and 5 ms for all time constants. The covariance matrix was initially diagonal with the standard deviation set to 0.003 and 3 ms for rule coefficients and time constants, respectively. We found that initializing the covariance matrix with small values relative to the search space led to fewer simulations in which weight growth was unbounded. The search space for rule coefficients and time constants was bounded to $[-10, 10]$ and $[0.05, 40]$ ms, respectively. The population size for each epoch of training was typically 14.

Each evaluation of a candidate learning rules was performed using the aggregate loss from 10 randomly initialized networks (see Supp. Section 1.1) to encourage CMA-ES to find plasticity rules that gracefully generalized to unobserved initial network structures and inputs. Importantly, the seeds of these simulation were held fixed from trial to trial; in a separate set of experiments, we attempted to draw a fresh set of networks and inputs for evaluation of the loss function, but found this drastically slowed learning.

#### 4 Comparison to existing model of sequence formulation

We compared discovered rules for sequence generation to a prior plasticity rule introduced by Fiete et al. [4]. The rule relies on spike timing dependent plasticity and bounds on both individual synapses as well as bounds on the total synaptic strength onto and out of a neuron. While the rule was originally formulated for both binary neurons and spiking networks, we adapt it here for continuous firing rate units. Under this rule, synapses evolve according to

$$\dot{w}_{ij} = \alpha w_{ij}(\tilde{x}_i x_j - x_i \tilde{x}_j) - \alpha\beta \left( \left[ \sum_k w_{kj} - W_{\max} \right]^+ + \left[ \sum_{k'} w_{ik'} - W_{\max} \right]^+ \right), \tag{8}$$

where all $w_{ij}$ are additionally bounded by some $w_{\max} < W_{\max}$. The first term in this rule implements a multiplicative (in that potentiation and depression are proportional to the synapse size) spike timing-dependence plasticity kernel (or the continuous firing rate analog). The second term bounds the sum of all synapses onto a neuron and out of a neuron to $W_{\max}$. Fiete et al. [4] demonstrated this rule generates wide synfire chains if groups of $N$ neurons receive correlated noise inputs and $w_{\max} = W_{\max}/N$.

To compare Eq. 8 to the discovered rule, we first optimized $(\alpha, \beta, w_{\max}, W_{\max})$ to minimize the decoder error (first term of Eq. 2 in the main text) in a setup identical to that described in Sec. 1.1. Once optimized, we used discovered rules trained with and without synaptic turnover and Eq. 8 to organize sequence generating networks by activating networks 400 times; 6 activations from the end of this period were used to train a linear decoder without regularization that attempted to decode time

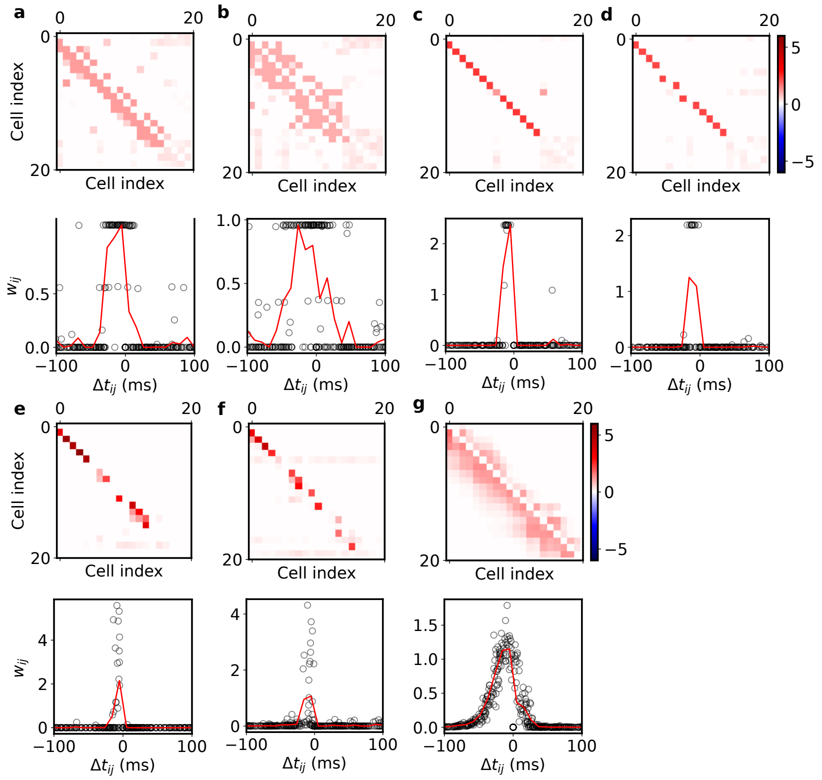


Figure 2: **Discovered rule organized dense feedforward structures**
Typical connectivity for sequences grown with different selections of Hebbian learning and heterosynaptic bound. **(a)** Connectivity matrix organized by additive Hebbian learning, summed synaptic bound, and a single synapse bound (top). Weight as a function of delay in firing time for all synapses in the network overlaid (bottom). Note: most nonzero synapses are at single synapse bound. **(b)** Same as (a), but with multiplicative Hebbian learning. **(c)** Same as (a), but with single synapse bound removed. Note: sequences become 1D. **(d)** Same as (c), but with single synapse bound removed. Sequences become 1D. **(e)** Connectivity organized by equation 10. **(f)** Connectivity organized when both Hebbian learning and firing rate bound are multiplicative. **(g)** Connectivity organized discovered rule, Eq. (6). Connectivity is dense and weight size depends on the relative time delay between firing times.

from the neural dynamics at 500 time points throughout each activation. Following this, networks underwent synaptic turnover for an additional 150 activations; synaptic turnover then ceased, and networks were given an additional 50 activations, after which the trained decoder's performance was evaluated on 200 time points from an additional 6 activations of the network.

#### 5 Learned rule generates dense feedforward connectivity relative to alternatives

Temporally asymmetric Hebbian learning and a mechanism that induces competition between synapses that drive the same postsynaptic target is a common recipe for sequence organization [4, 6–8]. Meta-learning selected a particular form of these two components both when synaptic turnover was included in training and when it was not. This led us to study the connectivity of sequences organized under different possible choices of these components (asymmetric Hebbian learning and synaptic competition) and compare them to those of the meta-learned rule. We fit the following learning rules using CMA-ES. In general, all coefficients and time constants were free parameters tuned by the model to produce the best encoding of time. The rules were

$$\dot{w}_{ij} = \tilde{x}_i x_j - x_i \tilde{x}_j - \beta \left( \left[ \sum_k w_{kj} - W_{\max} \right]^+ + \left[ \sum_{k'} w_{ik'} - W_{\max} \right]^+ \right), \tag{9}$$

$$\dot{w}_{ij} = w_{ij}(\tilde{x}_i x_j - x_i \tilde{x}_j) - \beta \left( \left[ \sum_k w_{kj} - W_{\max} \right]^+ + \left[ \sum_{k'} w_{ik'} - W_{\max} \right]^+ \right), \tag{10}$$

$$\dot{w}_{ij} = \tilde{x}_i x_j - x_i \tilde{x}_j - \tilde{x}_j x_j, \tag{11}$$

$$\dot{w}_{ij} = w_{ij}(\tilde{x}_i x_j - x_i \tilde{x}_j) - w_{ij} \tilde{x}_j x_j, \tag{12}$$

and

$$\dot{w}_{ij} = \tilde{x}_i x_j - x_i \tilde{x}_j - w_{ij} \tilde{x}_j x_j. \tag{13}$$

We fit Eqs. 9 and 10 both with and without a single synapse bound. The results are shown Fig. 2. We found that Eqs. 9 and 10 only grew dense feed-forward structures when a single synapse bound was imposed (Fig. 2, a-b). Removal of this bound generated 1D chains (Fig. 2, c-d). Eqs. 11 and 12 were generally less successful in organizing sequential activity, but grew structures where neurons received sparse synapses.

To explain these connectivity patterns, we considered the fixed points of individual synapses within them operating under the relevant rules. We first consider the effective learned rule of the meta-learned plasticity, Eq. 13. For $w_{ij}$ to be fixed, we require $\langle \dot{w}_i \rangle = 0$. Assuming synaptic evolution is slow, we find

$$\langle w_{ij} \rangle = \frac{\langle \tilde{x}_i x_j \rangle - \langle x_i \tilde{x}_j \rangle}{\langle \tilde{x}_j x_j \rangle}. \tag{14}$$

The synapse size is directly proportional to average potentiation under the Hebbian learning, as in Oja's rule, leading to connectivity shown in Fig. 2g.

We now consider what occurs if the postsynaptic bound does not scale with the synapse size, i.e. Eq. 11. Here, the potentiation under Hebbian learning must be equal on average to the postsynaptic activity bound, which is identical for all synapses that drive $j$. Thus the competition between synapses is winner-take-all, unless multiple synapses receive identical potentiation. We speculate that there are in general few presynaptic firing times relative to the postsynaptic firing times that receive equal potentiation under the first two terms of Eq. 11, thus sequences formed by this rule tend to be sparse (Fig 2c). The same argument applies to Eq. 9 when the single synapse bound is not imposed and for Eq. 12: assuming the synapse is nonzero, 12 may be rewritten as

$$\frac{d \log(w_{ij})}{dt} = \tilde{x}_i x_j - x_i \tilde{x}_j - \tilde{x}_j x_j, \tag{15}$$

which, after setting the right side to zero, is identical to the preceding case. In summary, that the last term in Eq. 13 is of greater order in the synapse size than the first two terms is important for generating dense connectivity without the use of single synapse bounds.

### 6 Meta-learning plasticity on all synapses

For completeness, we include the results of term-sensitivity analyses performed on solutions learned on unperturbed networks, networks with E→E turnover, and I→E turnover (Fig. 6). Note that E→E plasticity was largely similar to rules trained only on E→E synapses.

Formally, synapses of projection group $G$ evolved according to

$$\dot{w}_{ij}^{(G)} = \Theta(|w_{ij}^{(G)}|) \sum_k c_k^{(G)} F_k^{(G)}(x_i, x_j, w_{ij}^{(G)}, \tau_k^{(G)}), \tag{16}$$

where $F_k^{(G)}$ is the $k^{th}$ term of the plasticity rule acting on group $G \in \{\text{E}\to\text{E}, \text{E}\to\text{I}, \text{I}\to\text{E}\}$. $i$ ($j$) indexes the presynaptic (postsynaptic) neurons of $G$.

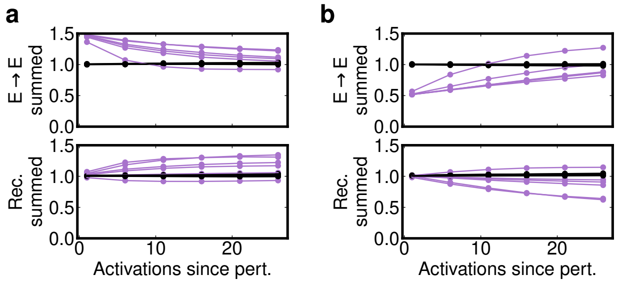


Figure 3: **Synaptic changes under rules learned with E→E turnover**
**(a)** Response of the magnitude of summed E→E weights $|W_i^{\rm exc}| = |\sum_k w_{k,i}^{E\to E}|$ (top) and summed recurrent weight $|W_i^{\rm rec}| = |\sum_k w_{i,k}^{\rm E\to I} w_{k,i}^{\rm I\to E}|$ (bottom) to the imposed scaling up of E→ E weights to a single cell. Values for neuron with perturbed inputs shown in light purple; all others black. Each line represents average values for a single learned rule over N=20 networks. **(b)** Same as (a), for imposed scaling down of E→ E weights to a single cell.

### 7 Single neuron perturbation experiments in networks with plasticity on all synapses

We performed three manipulations on 20 test networks under twelve different learned plasticity rules: six learned under E→E synaptic turnover and six under I→E synaptic turnover. In each manipulation, only the afferent synapses or inputs to a single neuron were modified. The targeted cell was always excitatory and randomly chosen. The three manipulations were:

1. Scaling down the excitatory afferents to an E cell by 50%.
2. Scaling up the excitatory afferents to an E cell by 50%.
3. Scaling up the stochastic input to a single E cell by a factor 10 ($\lambda^*_{\rm stoch} = 10\,\lambda_{\rm stoch}$).

In each case, plasticity rules were applied to initially random networks (see Section 1) for 400 network activations, after which the target neuron was manipulated and the synaptic change effected by the plasticity rules was measured at six subsequent time points, each spaced five activations apart.

We characterized the activity and synaptic connectivity of targeted cells. In particular, we recorded the sum of all excitatory weights onto the target cell as well as the strength of the recurrent inhibition, computed as

$$|W_i^{\rm rec}| = |\sum_k w_{i,k}^{\rm E\to I} w_{k,i}^{\rm I\to E}|. \tag{17}$$

In Fig. 6 and Supp. Fig. 3, we report these connectivity values normalized by pre-perturbation network averages across all neurons.

We additionally recorded changes in activity that stemmed from these manipulations and changes plasticity made to the network. We computed the peak amplitudes, durations, and timing of target cell firing. We defined the duration as $\int x_i(t)dt/x_i^{\rm (peak)}$ and the timing as $\int tx_i(t)dt/\int x_i(t)dt$. In Fig. 6d-f, Supp. Figs. 4a, c and 5a, c, we report peak amplitudes and durations normalized by pre-perturbation averages of the targeted cell. We report firing times relative to a pre-perturbation average (Supp. Figs. 4b, d and 5b, d).

We found that rules learned under different synpatic turnover settings and within the same setting largely converged in terms of function. Nearly all sets of rules increased (decreased) excitatory afferents onto the targeted cell and decreased (increased) recurrent inhibition in response to imposed scaling down (up) of excitatory afferents to the targeted cell (Fig 6a-c, Supp. Fig. 3). Plasticity rules acted to restore the peak amplitude, duration, and timing of an E cell's pre-perturbation activations (Supp. Figs. 4 and 5).

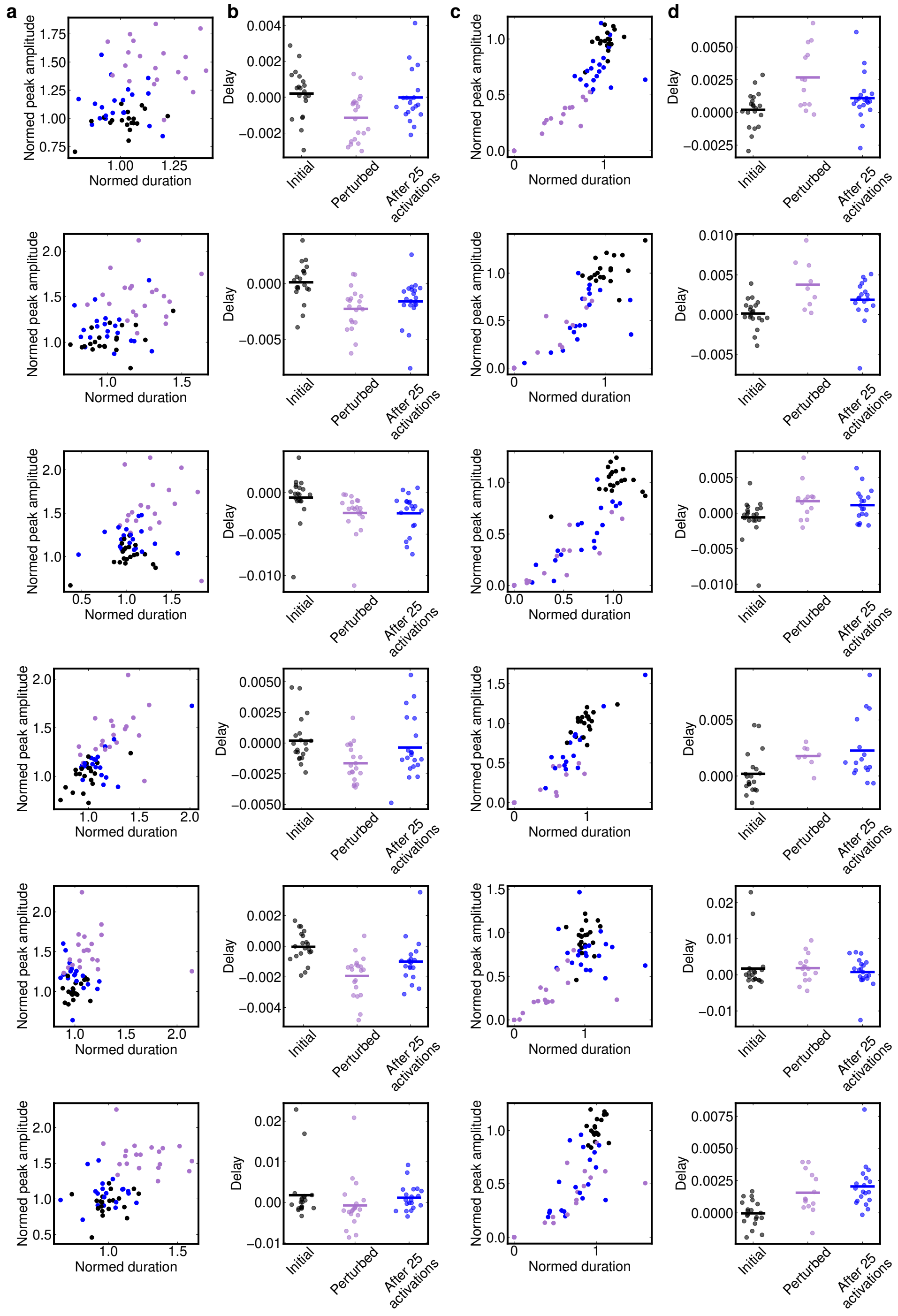


Figure 4: **Network response to single cell manipulations under all rules learned with E→E turnover (a)** Evolution of E cell postsynaptic firing patterns in the space of normalized peak amplitude and duration for six learned plasticity rules when a targeted E cell's afferent synapses are scaled up by 50%. Black points represent pre-perturbation values, light purple represent immediately post-perturbation, and blue represent 25 activations after perturbation. **(b)** Analysis of the timing of targeted E cell firing under the same rules and perturbations as (a). Note delays are relative to a pre-perturbation activation not pictured. Scaling up the excitatory afferents of the E cell initially shifts the firing time forward, but plasticity restores the original firing time of the E cell. **(c)** Same as (a), but with the excitatory afferents to a targeted E cell scaled down by 50%. Plasticity rules restore the initial peak amplitude and duration of targeted E cell responses. **(d)** Same as (b) when E cell afferents are scaled down by 50%. Perturbations initially delay firing of targeted cells, but four of six learned rules partially restore original firing time of targeted cells.

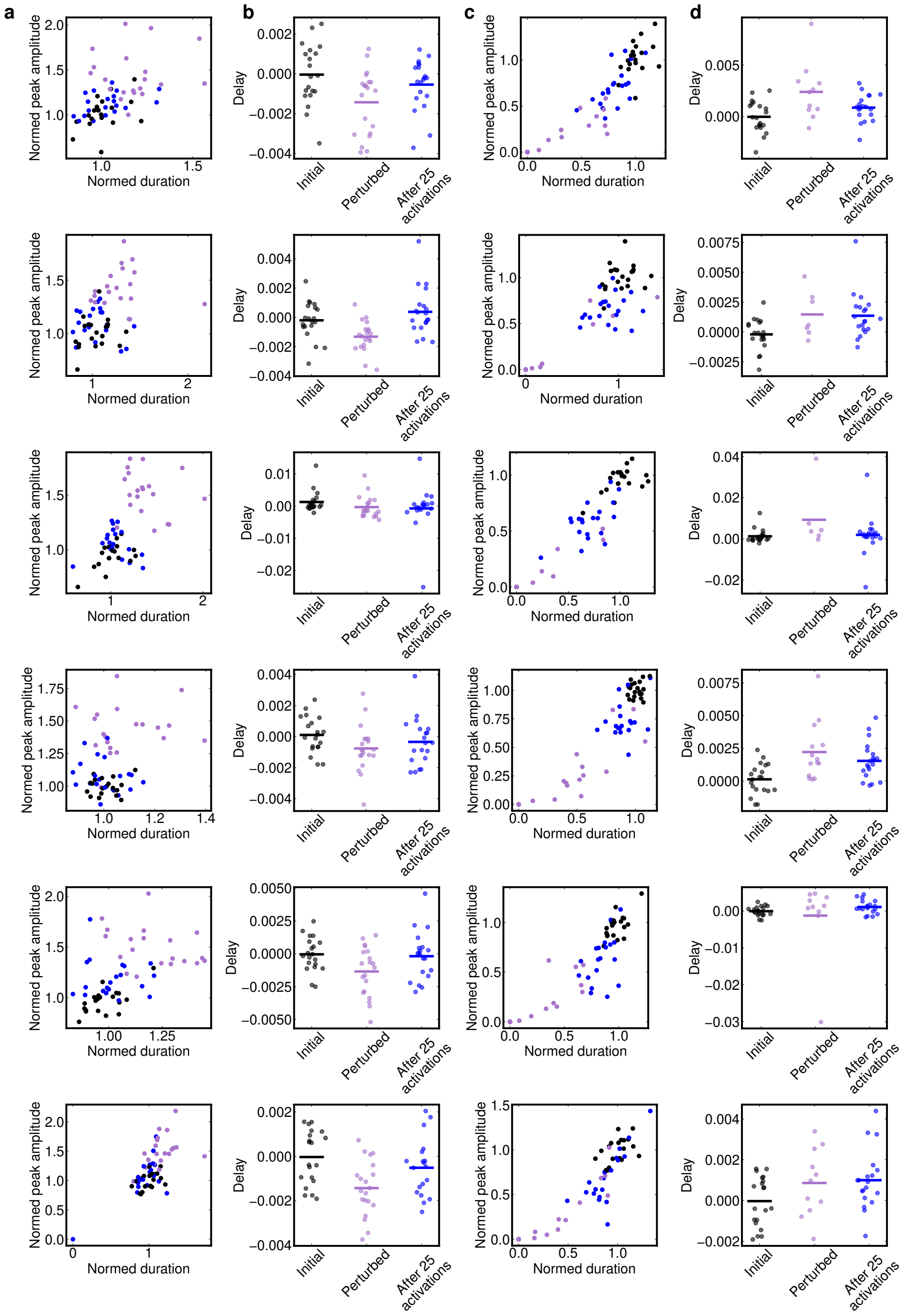


Figure 5: **Network response to single cell manipulations under all rules learned with I→E turnover** Identical to Supp. Fig. 4 (see caption) but for rules learned under I→E turnover.

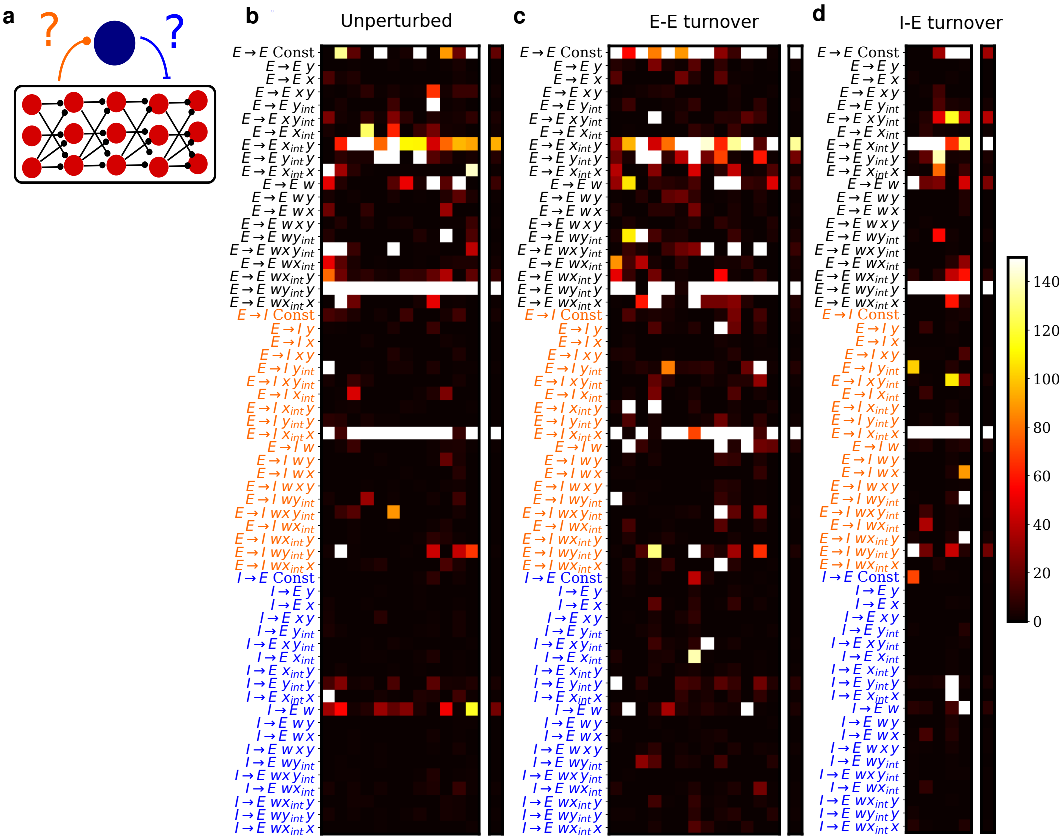


Figure 6: **Learning plasticity on all synapses**
**(a)** Schematic of synapses upon which additional plasticity will be learned. In previous simulations, these connections were held static. **(b)** Difference in medians between distribution of losses of full solutions run on 100 test networks and distribution of losses when one term is dropped on the same test set for solutions learned in the absence of synaptic turnover (left), **(c)** with turnover on E→E synapses, and **(d)** turnover on I→E synapses. Color of label indicates the synapses upon which each term acts. Isolated columns denote median loss for each term and type of perturbation. Note, temporally asymmetric Hebbian learning and heterosynaptic activity bound on E→E synapses remain amongst the most important terms by this measure across all types of perturbation. A presynaptic term on E→I synapses is also consistently important.

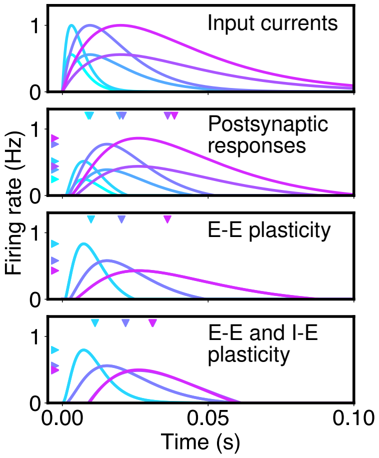


Figure 7: **Two plasticity mechanisms can correct timing**
Two plasticity mechanisms can better restore timing and postsynaptic firing pattern under a variety of inputs. Top panel: inputs to reduced neuron model described in Supp. Sec. 8. Second panel: initial firing rate responses of neuron to different inputs (colors matched to input). Third panel: firing rate responses of neuron evolving under E plasticity alone (see Supp. Sec. 8). Downward-pointing arrows indicate mean firing time of response and leftward-point arrows indicated peak firing rate. Bottom panel: firing rate responses of neuron evolving under E plasticity and I plasticity. Note mean firing times of responses and peak rates are more tightly clustered relative to those of responses of neuron with E plasticity alone.

#### 8 Reduced model of dual homeostatic control of E cell responses

We posit that the mechanism behind the improved robustness of the timing representation in networks with excitatory and inhibitory plasticity is compound homeostatic regulation of E cell postsynaptic response on different sets of synapses. To investigate this, we construct a model of a single neuron responding to a spectrum of excitatory inputs and investigate the stability of its postsynaptic response under different plasticity rules. Specifically, we compute the mean time relative to input onset and the magnitude of response under three different plasticity schemes:

1. Homeostatic conservation of squared magnitude of response on excitatory afferent
2. Homeostatic conservation of squared magnitude on excitatory afferent; conservation of magnitude on inhibitory afferent

We model the neuron as threshold linear unit, whose response is

$$r_i(t) = \left[ w_i \int_0^t e^{-(t-t')/\tau_{\text{in}}} r_{\text{in}}(t')dt' - b_i \right]^+ \tag{18}$$

Here, $w_i$ is the feed-forward excitatory weight, $b_i$ is the bias after $i$ inputs, and $\tau_{\text{in}}$ is the membrane time constant of the neuron. We assume that inhibition to the neuron is relatively tonic and is incorporated into $b_i$. We allow $w_i$ and $b_i$ to adjust after each input according to

$$w_{i+1} = w_i + \gamma_w \left( w_0 - \int_0^\infty r_i^2(t)dt \right) \tag{19}$$

and

$$b_{i+1} = b_i + \gamma_b \left( \int_0^\infty r_i(t)dt - b_0 \right), \tag{20}$$

where $w_0$, $\gamma_w$, $b_0$ and $\gamma_b$ are non-negative homeostatic set points and rates for the feed-forward weight and bias term, respectively. We drove the neuron with an alpha function,

$$r_{\text{in}}(t) = e a_{\text{in}}(t/\tau_{\text{in}})e^{-t/\tau_{\text{in}}}, \tag{21}$$

and tracked the evolution of the postsynaptic response in the phase space of peak response amplitude and response duration. We found a suitable choice of model parameters created a stable fixed point in this space. We further computed the delay between the average firing time of the response, $\int t r_i(t)dt / \int r_i(t)dt$, and the onset of input after 200 activations and found that plasticity tended to constrain the range of possible delays (Fig. 6j, right). We compared these dynamics to those of a neuron with fixed bias, and found it did not conserve the duration nor delay of the postsynaptic response (Fig. 6j).

In Supp. Fig. 7, we demonstrate how plasticity on I→E synapses in tandem with E→E plasticity improves timing robustness. A key aspect of this model is that it assumes inhibition is relatively constant while excitation is transient. Thus, I→E plasticity permits an E cell to adjust its resting potential in a manner that shortens or lengthens its postsynaptic response. Input that are too short (long) relative to the homeostatic setpoint are lengthened by decreasing (increasing) inhibition. E→E plasticity can then rescale the amplitude of the response to match the desired setpoint.

#### 9 Code availability

Code is available at on Github at `https://github.com/davidgbe/discovering_plasticity_organizing_circuits`.

#### References


[1] Anne Auger and Nikolaus Hansen. A restart cma evolution strategy with increasing population size. volume 2, pages 1769–1776, 01 2005. doi: 10.1109/CEC.2005.1554902.

[2] Y. Bengio, S. Bengio, and J. Cloutier. Learning a synaptic learning rule. ii:969 vol.2–, 1991. doi: 10.1109/IJCNN.1991.155621.

[3] Basile Confavreux, Friedemann Zenke, Everton J. Agnes, Timothy Lillicrap, and Tim P. Vogels. A meta-learning approach to (re)discover plasticity rules that carve a desired function into a neural network. In *34th Conference on Neural Information Processing Systems*, Vancouver, Canada, 2020.

[4] IR Fiete, W Senn, CZ Wang, and RH Hahnloser. Spike-time-dependent plasticity and heterosynaptic competition organize networks to produce long scale-free sequences of neural activity. *Neuron*, 65:563–76, 2010. doi: 10.1016/j.neuron.2010.02.003.

[5] J Jun and D Jin. Development of Neural Circuitry for Precise Temporal Sequences through Spontaneous Activity, Axon Remodeling, and Synaptic Plasticity. *PLOS ONE*, 2(8):e723, August 2007. ISSN 1932-6203. doi: 10.1371/journal.pone.0000723. URL `https://journals.plos.org/plosone/article?id=10.1371/journal.pone.0000723`. Publisher: Public Library of Science.

[6] U. Pereira and N. Brunel. Unsupervised learning of persistent and sequential activity. *Front Comput Neurosci*, 13:97, 2019. ISSN 1662-5188 (Print) 1662-5188. doi: 10.3389/fncom.2019.00097.

[7] Yevhen Tupikov and Dezhe Z Jin. Addition of new neurons and the emergence of a local neural circuit for precise timing. page 43, 2021.

[8] P. Zheng and J. Triesch. Robust development of synfire chains from multiple plasticity mechanisms. *Front Comput Neurosci*, 8:66, 2014. ISSN 1662-5188 (Print) 1662-5188. doi: 10.3389/fncom.2014.00066.
